# Significance tests for *R*^2^ of out-of-sample prediction using polygenic scores

**DOI:** 10.1101/2022.06.08.495250

**Authors:** Md. Moksedul Momin, Soohyun Lee, Naomi R Wray, S. Hong Lee

## Abstract

The coefficient of determination (*R*^2^) is a well-established measure to indicate the predictive ability of polygenic scores (PGS). However, the sampling variance of *R*^2^ is rarely considered so that 95% confidence intervals (CI) are not usually reported. Moreover, when comparisons are made between PGS based on different discovery samples, the sampling covariance of *R*^2^ is necessary to test the difference between them. Here, we show how to estimate the variance and covariance of *R*^2^ values to assess the 95% CI and p-value of the *R*^2^ difference. We apply this approach to real data to predict into 28,880 European participants using UK Biobank (UKBB) and Biobank Japan (BBJ) GWAS summary statistics for cholesterol and BMI. We quantify the significantly higher predictive ability of UKBB PGS compared to BBJ PGS (p-value 7.6e-31 for cholesterol and 1.4e-50 for BMI). A joint model of UKBB and BBJ PGS significantly improves the predictive ability, compared to a model of UKBB PGS only (p-value 3.5e-05 for cholesterol and 1.3e-28 for BMI). The proposed approach can also be applied to testing a significant difference between *R*^2^ values across different p-value thresholds. We also show that the predictive ability of regulatory SNPs is significantly enriched than non-regulatory SNPs for cholesterol (p-value 2.6e-19 for UKBB and 8.7e-08 for BBJ). We suggest that the proposed approach (available in R package ‘r2redux’) should be used to test the statistical significance of difference between pairs of PGS, which may help to draw a correct conclusion about the predictive ability of PGS.

## Introduction

Complex traits are affected by many risk factors including polygenic effects ^1–3^. Genetic profile analysis can quantify how polygenic effects are associated with future disease risk at the individual and population levels^4; 5^. Genetic profiling has potential benefits that can help people make informed decisions when they manage their health and medical care ^6–8^.

Genome-wide association studies (GWAS) have provided an opportunity to estimate genetic profile or polygenic scores (PGS) that can make individual risk predictions from genetic data ^4; 9–14^ Typically, the effects of genome-wide single nucleotide polymorphisms (SNPs) associated with complex traits are estimated in a discovery dataset, which are projected in an independent target dataset. Then, for each individual in the target samples the weighted genotypic coefficients according to the projected SNP effects (i.e. PGS) are derived and correlated with outcome (trait including affected/unaffected for disease) to quantify the prediction accuracy. The squared correlation or coefficient of determination (*R*^2^) have been a useful measure to quantify the reliability of PGS. Note that *R*^2^ is equivalent to squared regression coefficients if the dependent and explanatory variables are column-standardised ^15^.

Previously, we introduced a novel measure of *R*^2^ on the liability scale that can be comparable across different models and scales ^16^ when using disease traits or ascertained case-control data. Choi et al. ^12^ reported that this *R*^2^ measure on the liability scale outperforms the widely used Nagelkerke pseudo *R*^2^ in controlling for bias due to ascertained case-control samples. Nagelkerke pseudo *R*^2^ is not independent of the proportion of cases in the sample. In contrast, *R*^2^ on the liability scale is not dependent on the proportion of cases in the sample, but does require an estimate of the lifetime population prevalence of diseases.

Wand et al. ^11^ suggested that any PGS study should report *R*^2^ as an indicator of the predictive ability. Choi et al. ^12^ concluded that *R*^2^ is a useful metric to measure association and goodness of fit in the interpretation of PGS predictions. Many studies have demonstrated the predictive ability of PGS, using *R*^2^ ^12; 13; 17; 18^. However, the variance of *R*^2^ ^15^ has been rarely studied especially in the PGS analyses although it is the crucial parameter to estimate confidence intervals of *R*^2^. Furthermore, estimates of the covariance between a pair of *R*^2^ values (e.g., from two sets of PGS) are needed to assess if they are significantly different to each other, or if the ratio of two *R*^2^ values is significantly deviated from the expectation. This significance test for the difference or ratio is important when comparing two or multiple sets of PGS that are derived from different sets of SNPs, e.g., genomic partitioning or genome-wide association p-value thresholds (*p_T_*) analysis.

In this study, we use *R*^2^ measures and their variance-covariance matrix to assess if the predictive abilities of PGS based on different sources are significantly different to each other. We derive the variance and covariance of *R*^2^ values to generate estimates of its 95% CI and p-value of the *R*^2^ difference, considering two sets of dependent or independent PGS. We also derive the variance and covariance (i.e. information matrix) of squared regression coefficients in a multiple regression model, testing if the proportion of the squared regression coefficient attributable to SNPs in the regulatory region is significantly higher than expected (i.e. PGS-based genomic partitioning method). We apply this approach to real data to predict 28,880 European individuals using UK Biobank (UKBB) and Biobank Japan (BBJ) GWAS summary statistics for cholesterol and BMI.

## Methods

### PGS models

We use a linear model that regresses the observed phenotypes on a single or multiple sets of PGS. It is assumed that the phenotypes are already adjusted for other non-genetic and environmental factors (e.g. demographic variables, ancestry principal components (PCs)). A PGS model can be written as

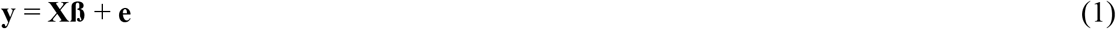

where **y** is the vector of standardised phenotypes of trait, **X** is a column-standardised N x M matrix including M sets of PGS, **ß** is the vector of regression coefficients and **e** is the vector of residuals. For example, with two sets of PGS (M=2), **X** and 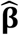 can be expressed as

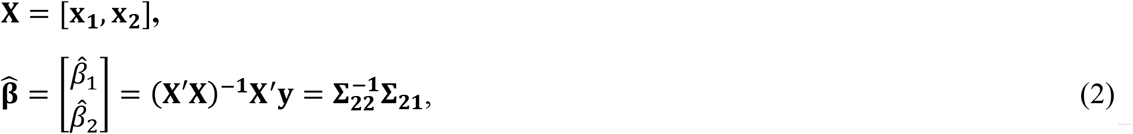

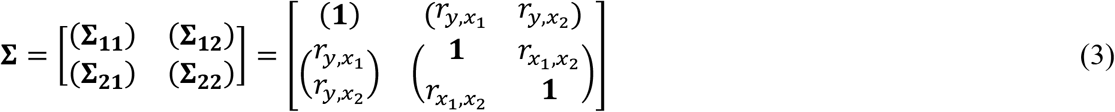

where *r_y,x_1__*, *r_y,x_2__* and *r_x_1_,x_2__* are correlations between **y** and the first PGS (**x_1_**), **y** and the second PGS (**x_2_**), and between the two PGS (**x_1_** and **x_2_**), respectively, in the sample. Using 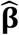 that are estimated in the multiple regression (eq. 2), the expected phenotypes (***ŷ***) can be obtained as

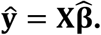

The coefficient of determination for this multiple regression model with **X** = [**x**_1_,**x**_2_] in eq. (1) can be written as

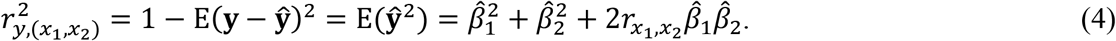

With a single set of PGS, i.e. M=1 and **X** = [**x**_1_] or [**x**_2_] in eq. (1), the expression of *R*^2^ can be reduced as

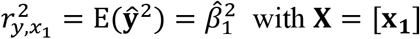

or

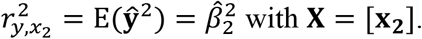

### Variance of *R*^2^

The distribution of *R*^2^ can be transformed to a non-central *χ^2^* distribution with mean = M+*λ* and variance = 2 × (*M* + 2*λ*) where 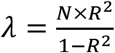 is the non-centrality parameter. For example, the variance of the transformed value for 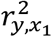 is

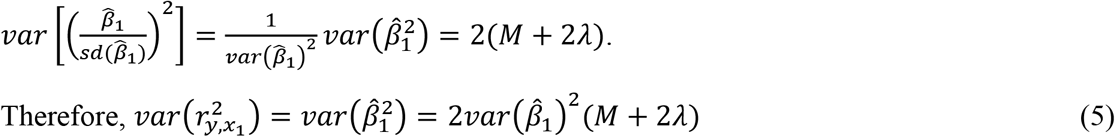

where 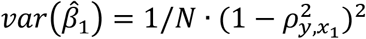 and 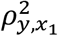 is the squared correlation in the population and can be approximated as 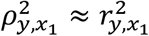 ^19; 20^.

In a similar manner, eq. (5) can be extended to multiple explanatory variables as

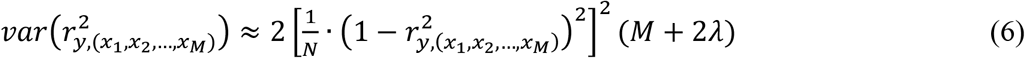

Wishart et al. ^21^ introduced a formula to obtain the variance of *R*^2^ as

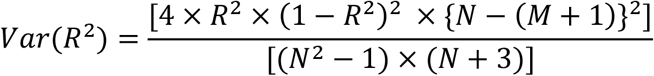

which provides an equivalent estimate as in eq. (6). The s.e. of *R*^2^ estimate is the square root of *var(R^2^*).

### Variance of the difference between two *R*^2^ values

Following Olkin and Finn ^15^, we use the delta method to estimate the variance of the difference between *R*^2^ values based on two sets of PGS (**x_1_** and **x_2_**). Assuming that the difference of *R*^2^ values can be formulated as a function of the correlations, i.e. *f*(*r*_*y,x*_1__, *r*_*y,x*_2__, *r*_*x*_1_,*x*_2__) the delta method approximates the variance of the difference as

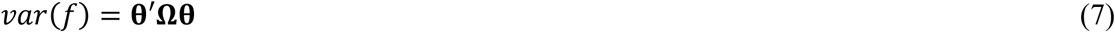

where 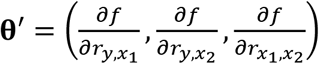 is the derivatives of *f* with respect to the correlations and

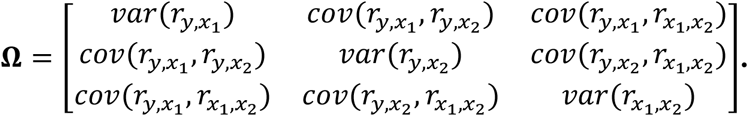

Each element of Ω is shown in Olkin and Finn ^15^ (also see Appendix).

From eq. (7), the following variances of differences can be estimated and used in our PGS analyses.

#### 1. *R*^2^ difference when using different discovery samples to generate the PGS

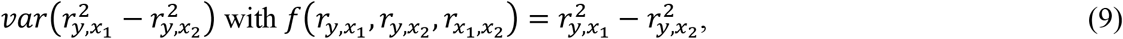

which allows us to compare two PGS models that are not nested to each other (see *‘R^2^* difference when using different information sources’ in Results section), for which the conventional log-likelihood ratio test cannot be applied.

In eq. (9), the values of 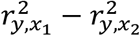 from random samples in the population are normally distributed when the sample size is sufficient ^15^. Assuming that our PGS analysis is sufficiently powered (N > 25,000), the p-value for the significance test of the difference can be derived from

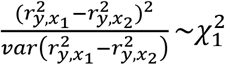

and the 95% confidence interval is

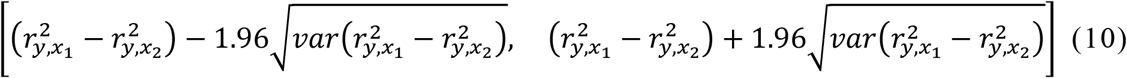

#### 2. *R*^2^ difference when using nested models

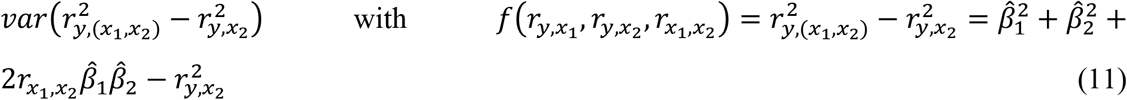

where 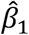 and 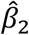 are the estimated regression coefficients from a multiple regression (eq. 2), calculated from **Σ** (see eq. 2–4). Again, the derivative with respect to each of the correlations can be obtained for this function (eq. 8). Note that the comparison between 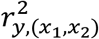 and 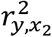 is equivalent to the log-likelihood ratio test (i.e. **y = x_1_***β*_1_ + **x_2_***β*_2_ **+ e** vs. **y = x_2_***β*_2_ **+ e**)^15^.

The values of 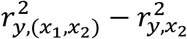 in eq. (11) from random samples in the population follows a non-central chi-squared distribution with a non-centrality parameter 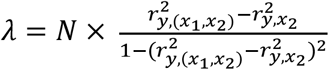. The p-value for the significance test of the difference can be derived from

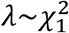

and the 95% confidence interval is

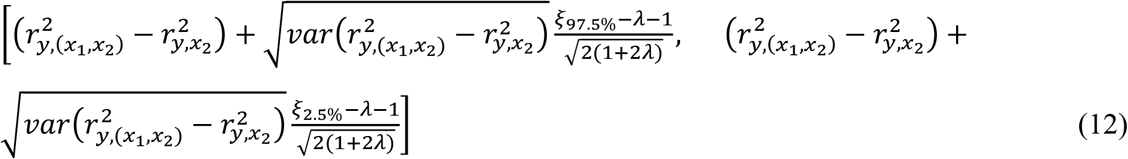

where *ξ*_%_ is the value at the percentile of the inverse of non-central chi-squared cumulative distribution function with mean = *λ* + 1 and d.f. = 1.

When the sample size is large, the values of 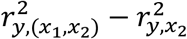 from random samples in the population are normally distributed ^15^. The p-value for the significance test of the difference can also be derived from

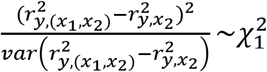

and the 95% confidence interval is

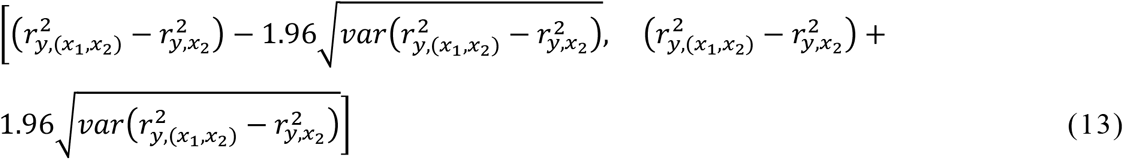

Note that eq. 12 and 13 are equivalent when the sample size is sufficient ^15^.

#### 3. *R*^2^ difference when using two independent sets of PGS

In this case, there is no correlation structure between two independent sets of PGS (*r*_*x*_1_,*x*_2__ = 0, e.g., male and female PGS), therefore, the variance of *R*^2^ difference is simply the sum of the variances of each *R*^2^ value, which can be obtained from eq. (5). For example, assuming *r*_x_1_,*x*_2__ = 0, the variance of *R*^2^ difference can be written as

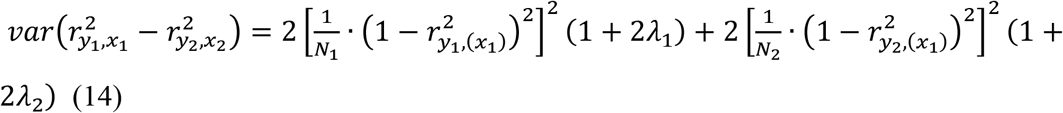

where *y*_1_ and *y*_2_ are the vectors of standardised phenotypes and *N*_1_ and *N*_2_ are the sample sizes for the two independent sets of PGS. The non-centrality parameters (*λ*_1_ and *λ*_2_) for two dependent PGS can be written as

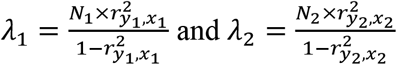

p-value for the significance test of the difference can be derived from 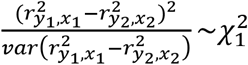 and the 95%confidence interval is

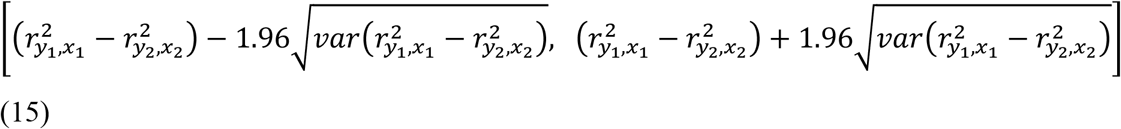

#### 4. PGS-based genomic partitioning analysis

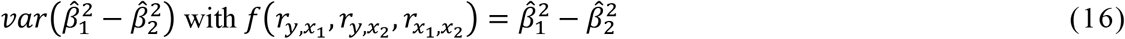

where 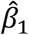 and 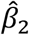 are the estimated regression coefficients from a multiple regression (eq. 2), calculated from **Σ** (eq. 3). Therefore, it is possible to get the derivative with respect to each of the correlations, *r*_*y,x*_1__, *r*_*y,x*_2__ and *r*_*x*_1_,*x*_2__ in eq. (8). 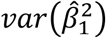 and 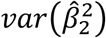 can be also obtained in a similar manner. Thus, we can get the variance covariance matrix 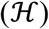, i.e. the information matrix, as

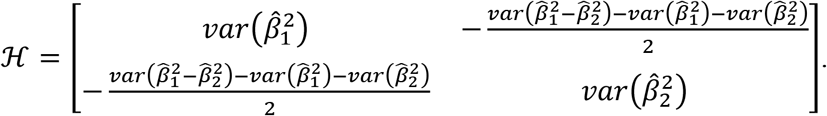

The 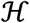 can be used to estimate the variance of the difference between 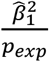 and 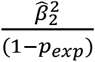 as

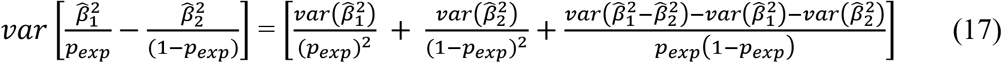

where the expected proportion of phenotypic variance explained by *x*_1_ (PGS1) can be calculated from prior information, referred to as *p_exp_* = # SNPs used for PGS1 / total # SNPs. This variance can be used to test if two squared regression coefficients scaled by their expectations are significantly different to each other.

Analogous to eq. (9), the values of 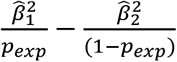 in eq. (17) with random samples in the population are asymptotically normal ^15^. Using a Wald test, the p-value for the significance test of the difference can be derived from

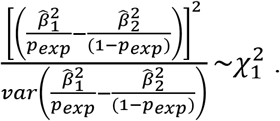

The 95% confidence interval of the ratio is

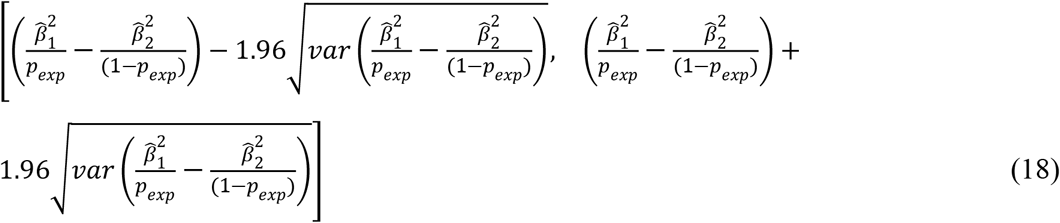

### Data

The UK Biobank is a large-scale biomedical database, that comprises 0.5 million individuals who had been recruited between 2006 to 2010 and their age ranged between 40 and 69 years ^22; 23^. The data consists of health-related information for samples who are genotyped for genome-wide SNPs. A stringent quality control (QC) process was applied to UKBB data that excludes individuals with non-white British ancestries, mismatched gender between reported and genotypic information, genotype call rate <0.95 and putative sex chromosome aneuploidy. The SNP QC criteria filtered out SNPs with an imputation reliability <0.6, missingness >0.05, minor allele frequency (MAF) <0.01, Hardy-Weinberg equilibrium p-value < 10^-07^. We also applied a relatedness cut-off QC (>0.05) so that there was no high pairwise relatedness among samples. After QC, 288,792 individuals and 7,701,772 SNPs were remained.

### Discovery GWAS data

Ninety percent of the individuals from the QCed data (N=288,792) were randomly selected as discovery samples (N=259,912, SNP # =7,701,772) to generate GWAS summary statistics (UKBB hereafter). For the GWAS with the 259,912 UKBB discovery samples, we used BMI and cholesterol that were adjusted for age, sex, birth year, Townsend deprivation Index (TDI), education, genotype measurement batch, assessment centre and the first 10 ancestry principal components (PCs) using a linear regression.

We also have access to Japanese Biobank (BBJ) (http://jenger.riken.jp/en/result) GWAS summary statistics (BBJ hereafter) for BMI ^24^ (N=158,284) and cholesterol ^25^ (N=128,305) for 5,961,601 SNPs.

### Target data

Ten percent of the individuals from the QCed data (N=288,792) were randomly selected as an independent target dataset (N=28,880 and SNP # =7,701,772) that were non-overlapping and unrelated with the UKBB and BBJ discovery samples. In the PGS analyses, we used only 4,113,630 SNPs that were common between UKBB and BBJ GWAS data after excluding ambiguous SNPs and SNPs with any strand issue.

In the target dataset (N=28,880), the phenotypes of each trait were adjusted for age, sex, birth year, TDI, education, genotype batch, assessment centre and the first 10 PCs using a linear regression. The pre-adjusted phenotypes were correlated with PGS estimated in the following step. For each trait, we used the UKBB and BBJ GWAS summary statistics to estimate two sets of PGS (UKBB PGS vs. BBJ PGS for the target individuals (n=28,880), using PLINK2 – score function ^26^. Then, we estimated the correlation between the PGS and pre-adjusted phenotypes to obtain *R*^2^ values in the PGS analyses.

### Functional annotation of the genome

We annotated the genome using pre-defined functional categories (regulatory vs. non-regulatory genomic regions) ^27^. Regulatory region includes SNPs from coding regions, untranslated regions (UTR) and promotors. Non-regulatory region includes all the other regions except the regulatory region. The number of SNPs belong to regulatory and non-regulatory is 158,653 and 3,954,947 (i.e. 4% of the total SNPs are located in the regulatory region).

### Simulation of dependent and explanatory variables

For a quantitative trait, we simulated dependent variable (*y*) and PGS (*x*_1_ and *x*_2_), varying the correlation structure of 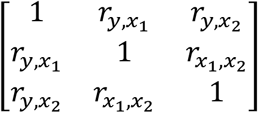 and the sample size (detailed simulation parameters are shown in Supplementary Figures 1-9). For a disease trait, the same simulation procedure was used, and the simulated quantitative phenotypes were transformed to binary responses using a liability threshold model with a population prevalence of k=0.05. For example, case-control status was assigned to individuals according to their standardised quantitative phenotypes (i.e. liability), i.e. cases have liability greater than a threshold such that the proportion of cases is k=0.05. The empirical variances of 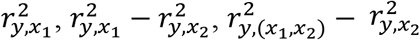 and 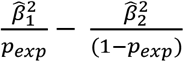 were obtained over 10,000 replicates, which were compared to the theoretical variances estimated using eqs. (6), (9), (11) and (17), respectively.

## Results

### Simulation verification

The theory of the proposed method has been explicitly verified using simulations, varying sample size and values of 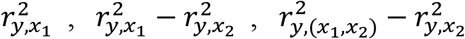 and 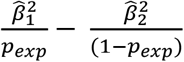 (**Supplementary Figures 1 – 9**). The empirical variances obtained from 10,000 simulated replicates are almost perfectly correlated with the theoretical variance for the values of, 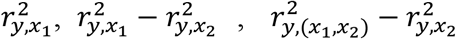 and 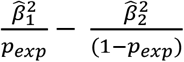 when varying the sample size (**Supplementary Figures 1 – 4**) and when varying *R*^2^ values (**Supplementary Figures 5 – 8**). When considering two independent PGS, the theoretical and empirical variances are also agreed well (**Supplementary Figure 9**).

### *R*^2^ difference when using different information sources (UKBB vs. BBJ)

It is of interest to determine whether different information sources (e.g., ancestries) have significantly different predictive abilities in PGS analyses, which can be assessed using eqs. (9) and (10). **Figure 1** illustrates that when predicting the 28,880 European target samples, the coefficient of determinations (*R*^2^) with the UKBB and BBJ PGS were 0.024 (95% CI = 0.02-10.028) and 0.003 (95% CI = 0.002 - 0.004), respectively, for cholesterol. However, these *R*^2^ values and CIs cannot be used to assess their difference because the two sets of PGS are not independent. Furthermore, the two PGS models with UKBB and BBJ are not nested to each other, therefore, the likelihood ratio test could not be used either. For this problem, we used eqs. (9) and (10) to obtain the variance, 95% CI (0.0247 - 0.0175) and p-value (7.6e-31) of the *R*^2^ difference, accounting for the dependency between UKBB and BBJ PGS, for cholesterol (**Figure 1**). Similarly, the test statistics of the *R*^2^ difference was obtained for BMI, 0.035 - 0.046 for 95% CI and p-value = 1.4e-50 (**Figure 1**).

**Figure 1:**
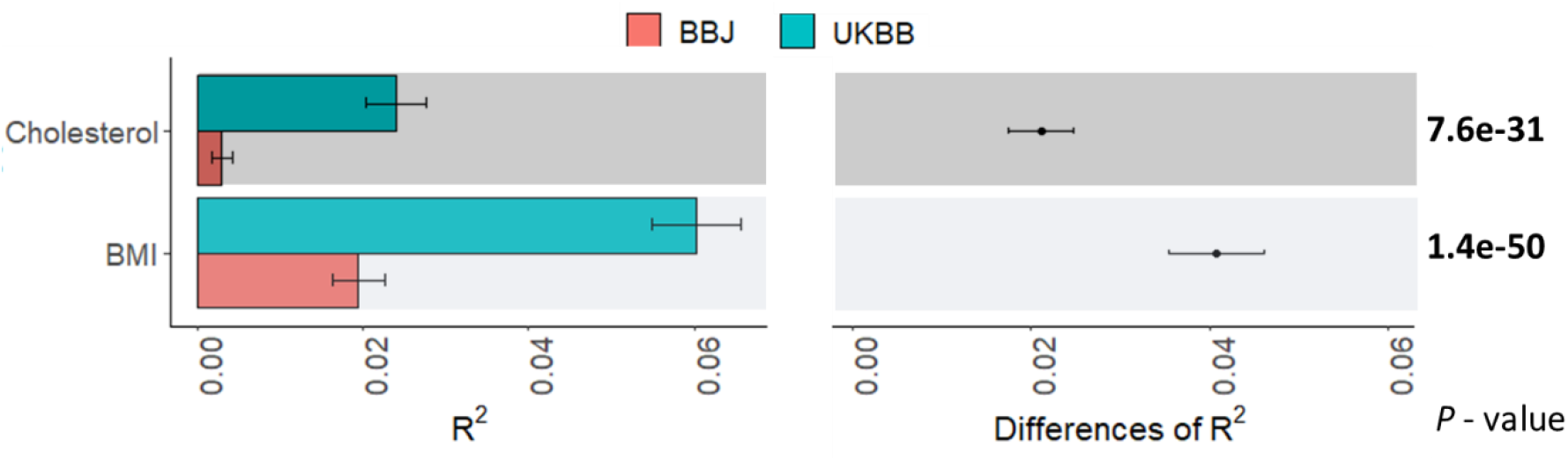
The predictive ability (*R*^2^) of PGS when predicting 28,880 European individuals using UKBB or BBJ discovery GWAS dataset. **Left panel:** The main bars represent *R*^2^ values and error bars correspond 95% confidence intervals. Two sets of GWAS summary statistics were obtained from UKBB and BBJ discovery GWAS datasets to estimate two sets of PGS. **Right panel:** Dot points represent the differences of *R*^2^ values between UKBB and BBJ PGS models, and error bars indicate 95% confidence intervals of the difference.

It is also interesting to test if BBJ PGS provides a significant improvement in the predictive ability, in addition to UKBB PGS, when predicting the 28,880 European target samples. **Figure 2** compares *R*^2^ value with each UKBB or BBJ PGS to *R*^2^ value from a joint model fitting UKBB and BBJ PGS simultaneously. Using eq. (11) and (12), we acquired the variance, 95% CI (0.0001–0.001) and p-value (3.5e-05) of *R*^2^ difference when comparing the joint model with a single model with UKBB, indicating that BBJ PGS contributed to a significant improvement for cholesterol. Similarly, BBJ PGS improved the predictive ability significantly (p-value = 1.3e-28) for BMI. As expected, excluding UKBB PGS from the joint model substantially decreased the prediction accuracy (p-value = 1.6e-136 for cholesterol and 3.0e-308 for BMI).

**Figure 2:**
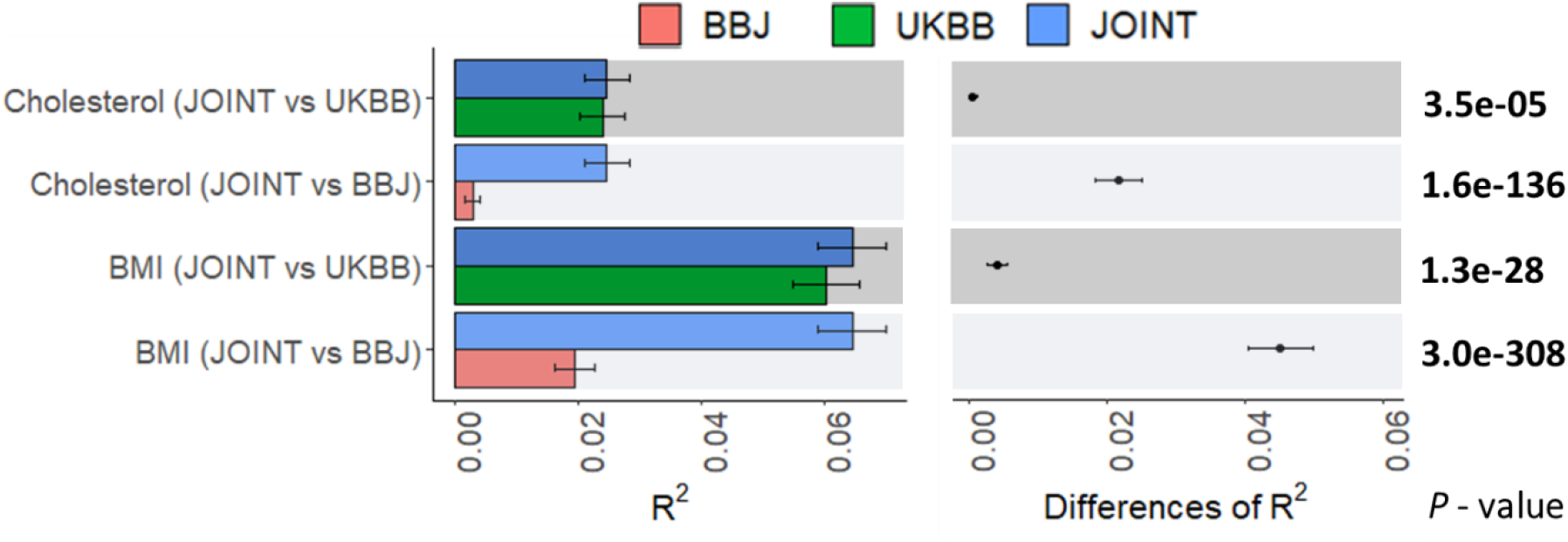
The predictive ability (*R*^2^) of the UKBB or BBJ PGS model or a joint model of UKBB and BBJ when predicting 28,880 European individuals. **Left panel:** The main bars represent *R*^2^ values and error bars correspond 95% confidence intervals. Two sets of GWAS summary statistics were obtained from UKBB and BBJ discovery GWAS datasets to estimate two sets of PGS, i.e. UKBB and BBJ PGS. In addition, a joint model fitting both UKBB and BBJ PGS was compared. **Right Panel:** Dot points represent the differences of *R*^2^ values between the joint model and UKBB or BBJ PGS model, and error bars indicate 95% confidence intervals of the difference.

### *R*^2^ difference when using two independent sets of PGS (male vs. female)

We were also interested in testing if the PGS could predict the adjusted phenotypes of the target individuals equally well for males and females. In this case, there is no correlation structure between male and female PGS, therefore, the variance of *R*^2^ difference is simply the sum of the variances of each *R*^2^ value, which can be obtained from eq. (5) or (6). **Supplementary Figure 10** shows that there was no significant difference between male and female PGS in their predictive ability for cholesterol and BMI whether using UKBB or BBJ discovery GWAS dataset.

### PGS with genome-wide association p-value thresholds (*p_T_*)

PGS also has been widely used to determine which *p_T_* provides the highest prediction accuracy, for example, using PGS software such as PLINK ^26; 28^. However, there is a lack of test statistics that can assess if the predictive ability of the best-performed *p_T_* is significantly different from the other *p_T_*. **Figure 3a** illustrates that *R*^2^ value is the highest at *p_T_=* 0.3 when predicting 28,880 European individuals in the target dataset, using BBJ discovery GWAS dataset (BMI). However, it is not clear if the predictive ability at *p_T_* = 0.3 is significantly higher than the adjacent *p_T_* (e.g. *p_T_* = 0.2 or 0.4), and it may be important to report *p_T_* of which the predictive ability is not statistically different from the best-performed *p_T_*. Using eqs. (9) and (10), we assessed the significance of difference between the best-found *p_T_* and each of the other *p_T_* (**Figure 3b**). From this analysis, we found that the best-performed *p_T_* was not significantly different from *p_T_* ranging between 0.1 – 1, but significantly different from *p_T_* ≤ 0.05 (**Figure 3b**). When using UKBB discovery GWAS dataset to predict the 28,880 European individuals, the highest *R*^2^ value at the *p_T_* of 1 was significantly different from all the other *p_T_*(**Supplementary Figure 11b**).

**Figure 3:**
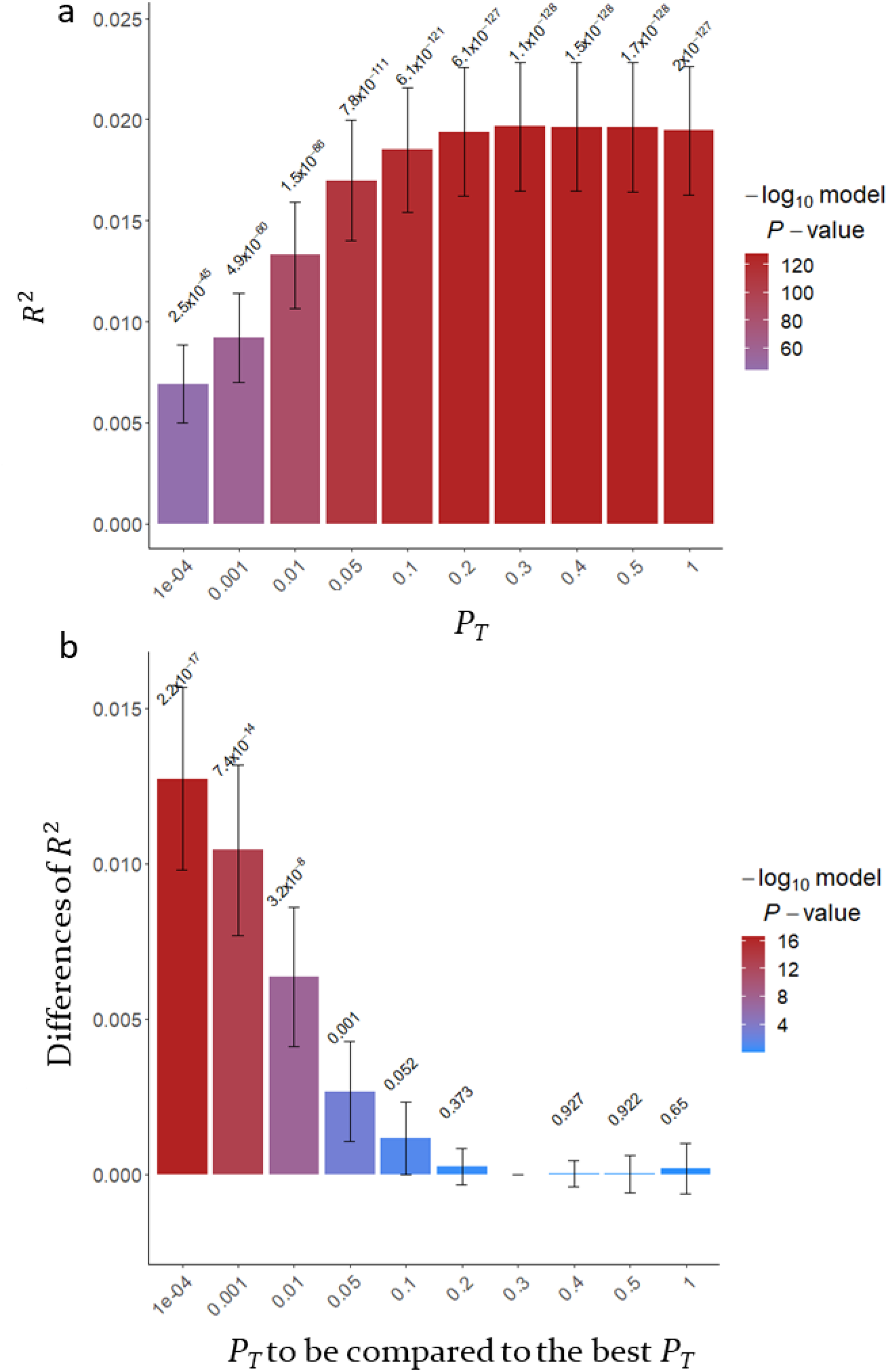
The predictive ability (*R*^2^) of PGS estimated based on SNPs below *p_T_* when predicting BMI in 28,880 European samples using BBJ discovery samples (GWAS summary statistics). a) The main bars represent *R*^2^ values and error bars correspond 95% confidence intervals. The values above 95% CIs are p-values indicating that *R*^2^ values are not different from zero. b) The main bars represent the difference of *R*^2^ values between the corresponding *p_T_* and the best-performed *p_T_* and error bars indicate 95% confidence intervals. The values above 95% CIs are p-values indicating the significances of differences between the pairs of *R*^2^ values.

Interestingly, the highest *R*^2^ value was found at *p_T_* = 1e-04 (**Figure 4a**) when predicting the European target samples using BBJ discovery GWAS dataset for cholesterol, which was not statistically different from *p_T_* = 0.001, but was significantly higher than the other *p_T_* (**Figure 4b**). For the same target samples and trait, the best *R*^2^ value was obtained from *p_T_* = 0.01 when using UKBB discovery GWAS dataset (**Supplementary Figure 12a**). Except for *p_T_* = 0.01, 0.05 and 0.1, *R*^2^ values at the other *p_T_* were significantly different from the best *R*^2^ values (**Supplementary Figure 12b**).

**Figure 4:**
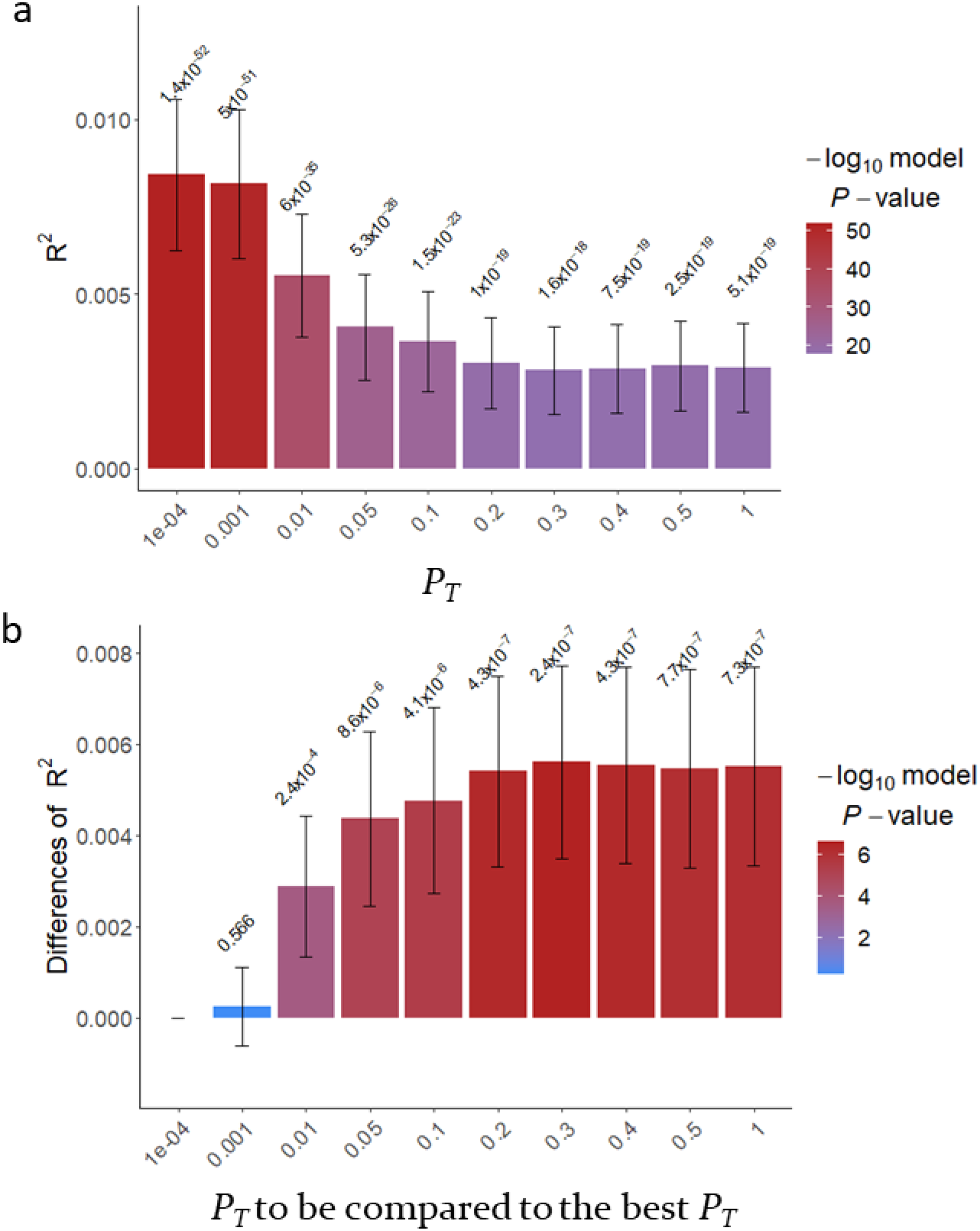
The predictive ability (*R*^2^) of PGS estimated based on SNPs below the *p_T_* when predicting cholesterol in 28,880 European samples using BBJ discovery samples (GWAS summary statistics). a) The main bars represent *R*^2^ values and error bars correspond 95% confidence intervals. The values above 95% CIs are p-values indicating that *R*^2^ values are not different from zero. b) The main bars represent the difference of *R*^2^ values between the corresponding *p_T_* and the best-performed *p_T_* and error bars indicate 95% confidence intervals. The values above 95% CIs are p-values indicating the significances of differences between the pairs of *R*^2^ values.

### PGS-based genomic partitioning analyses

Genomic partitioning analyses have been widely applied ^27; 29-31^. Such analysis could be useful in the PGS context. Using eq. (17) and the information matrix, 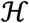, we can estimate the variance of the difference between, 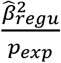 and 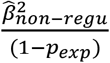 where 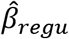 and 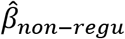 are the estimated regression coefficients from a multiple regression (eq. 2), and assess if the differences is significant (i.e. the coverage of the SNPs belonged to the category). For example, we partitioned the genome-wide SNPs into the regulatory (158,653) and non-regulatory regions

(3,954,947), following Gusev et al. ^27^, resulting 4% of SNP coverage for the regulatory region as the expectation. We simultaneously fit two sets of PGS from regulatory and non-regulatory to get 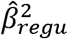 and 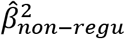, using a multiple regression, and assess if the difference, 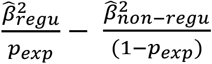, is significant (eq. 18). **Figure 5** shows that the predictive ability of regulatory SNPs was significantly higher than nonregulatory SNPs (p-value = 2.6e-19 for UKBB and 8.7e-08 for BBJ) for cholesterol. In contrast, the predictive ability of regulatory SNPs was not different from the expectation (p-value = 1.0e-01 for UKBB and 8.2e-01 for BBJ) for BMI.

**Figure 5:**
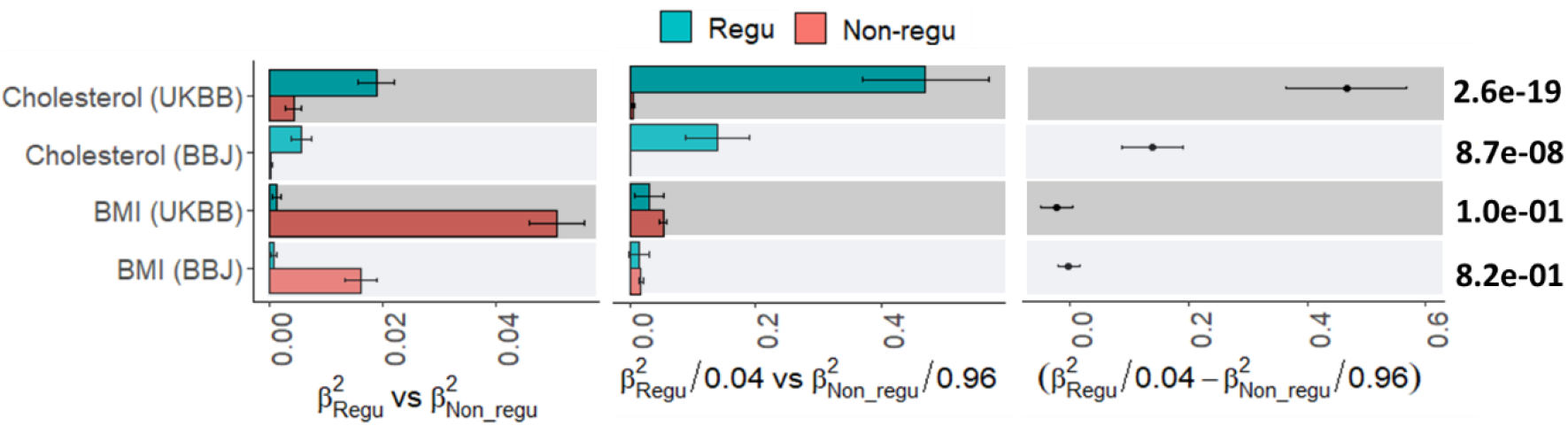
PGS-based genomic partitioning method to assess if the predictive ability is enriched in the regulatory region for cholesterol and BMI. Here 0.04 is the expectation for the regulatory SNPs based on the proportion of SNPs allocated to this annotation. **Left panel:** The main bars represent squared regression coefficients attributable to SNPs in the regulatory region 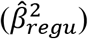 and non-regulatory region 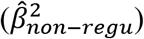, and error bars correspond 95% confidence intervals when predicting 28,880 European samples using UKBB or BBJ GWAS summary statistics. **Middle panel:** The main bars represent the ratio of 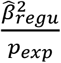 and 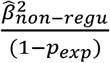 and error bars correspond 95% confidence intervals when predicting 28,880 European samples using UKBB or BBJ GWAS summary statistics. **Right panel:** Dot points represent the difference, 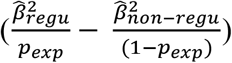 between regulatory region 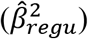 and non-regulatory region 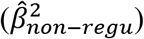 and error bars indicate 95% confidence intervals of the ratio differences.

### Application to binary responses and ascertained case-control data

The proposed method is also explicitly verified using simulation for binary or case-control data, varying sample size and values of 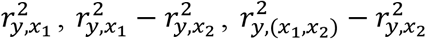 and 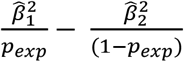 (**Supplementary Figures 13 – 20**). The empirical variances obtained from 10,000 simulated replicates are almost identical with the theoretical variances for the values of 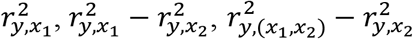 and 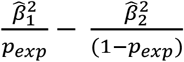 when varying the sample size (**Supplementary Figures 13 – 16**) and when varying *R*^2^ values (**Supplementary Figures 17 – 20**). In the case of ascertained case-control, a similar pattern is shown, i.e. the empirically observed variances obtained from 10,000 simulated replicates are agreed well with the theoretical variances for the values (**Supplementary Figures 21 – 24**). This finding shows that the proposed method can be applied to test the significance of difference between predictive abilities of PGS for binary traits and ascertained case-control traits when *R*^2^ is not very high (< 0.1). Note that the empirical and theoretical variances become disagreed when *R*^2^ values on the observed scale are more than 0.1 for binary responses and ascertained case control (**Supplementary Figures 25 and 26**). Although *R*^2^ value > 0.1 is not frequently observed in the current PGS studies (**Supplementary Table 2**), a careful interpretation is required for the variance of such high *R*^2^, and we would not recommend using the theoretical approximation.

## Discussion

*R*^2^ has been widely used to measure the predictive ability of PGS ^13^. However, the confidence interval of *R*^2^ has rarely been reported, and the test statistic for the difference of two *R*^2^ values has not been well documented. Here, we show how to get the variance of each estimated *R*^2^ value, and covariance between two *R*^2^ estimates (from two sets of PGS) that can be used to assess if they are significantly different to each other.

Martin et al. ^18^ reported that the PGS prediction accuracy is higher when discovery and target samples are from the same ancestry background, compared to when the samples are from different ancestries. However, they did not formally assess the statistical significance of the increase (no p-value provided). More importantly, they did not consider the correlation structure between predictors when they compared two PGS (in their Figure 4). We applied the proposed approach and found that the predictive ability of PGS based on UKBB discovery GWAS is significantly higher than that of PGS based on BBJ discovery GWAS, by formally deriving the 95% CI and p-value of the *R*^2^ difference.

Many studies evaluating PGS use the *p_T_* method ^12^, and report the *p_T_* that maximises performance. This provides useful information when inferring the genetic architecture of the trait of interest and when fine-tuning *p_T_* as a hyper-parameter in PGS methods ^28; 32–34^ For such cases, it may be crucial to determine if the best-performed *p_T_* is genuinely better than other (adjacent) *p_T_* or it occurs just by random chance (i.e. sampling error). Our proposed approach can formally assess statistical difference among *p_T_*, providing 95% CI of the difference with a significance p-value.

We also derived an information matrix of squared regression coefficients in a multiple regression model, establishing a PGS-based genomic partitioning method that could test if the ratio of two squared regression coefficients is significantly deviated from its expectation given the proportion of SNPs allocated to each partition. This is analogous to the existing genomic partitioning approaches, using GREML or LDSC ^27; 29–31^ that may have an overfitting issue because SNP effects and genomic partitioning are estimated in the same samples.

In conclusion, we show how to estimate the variance and covariance of *R*^2^ estimates to quantify the 95% CI and p-value of the difference and ratio when considering a pair of PGS, which is available in R package ‘r2redux’ (see Appendix B). We suggest that the proposed approach should be used to test the statistical significance of difference and ratio between pairs of PGS, which may help to draw a correct conclusion about the predictive ability of PGS.

## Code availability

The genotype and phenotype data of the UK Biobank can be accessed through procedures described on its webpage (https://www.ukbiobank.ac.uk/) and summary statistics of BMI and cholesterol from Japanese Biobank (BBJ) can be obtained from its website (http://jenger.riken.jp/en/result) PLINK2 version can be downloaded from https://www.cog-genomics.org/plink/r2redux can be downloaded from (https://github.com/mommy003/r2redux_version4) and to be added in the CRAN soon (also see Appendix B).

## Acknowledgements

This research is supported by the Australian Research Council (DP190100766). The R package development is supported by Cooperative Research Program for Agriculture Science and Technology Development (PJ0160992021) from the Rural Development Administration, Republic of Korea. We thank the staffs and samples of the UK Biobank and Biobank Japan for their important contributions. Our reference number approved by UK Biobank is 14575. The UK Biobank is funded by the UK Department of Health, the Medical Research Council, the Scottish Executive, and the Wellcome Trust medical research charity. The analyses were performed using computational resources provided by the Australian Government through Gadi under the National Computational Merit Allocation Scheme (NCMAS), and HPCs (Tango and Statgen server) managed by UniSA IT. NRW acknowledges funding from the NHMRC (1113400, 1173790).

## Author contribution

S.H.L. and N.R.W. conceived the idea. S.H.L. derived theory and supervised the study. M.M.M performed the analysis. M.M.M and S.H.L verified the theory and analytical methods, and made the R package, with support from S.L. S.H.L and M.M.M wrote the first draft of the manuscript. N.R.W. and S.L. provided critical feedback and suggestions. All the authors discussed the results and contributed to the final manuscript.

## Appendix A. The elements of Ω in eq. (7)

Following Olkin and Finn ^15^, each element of **Ω** in eq. (7) can be expressed as

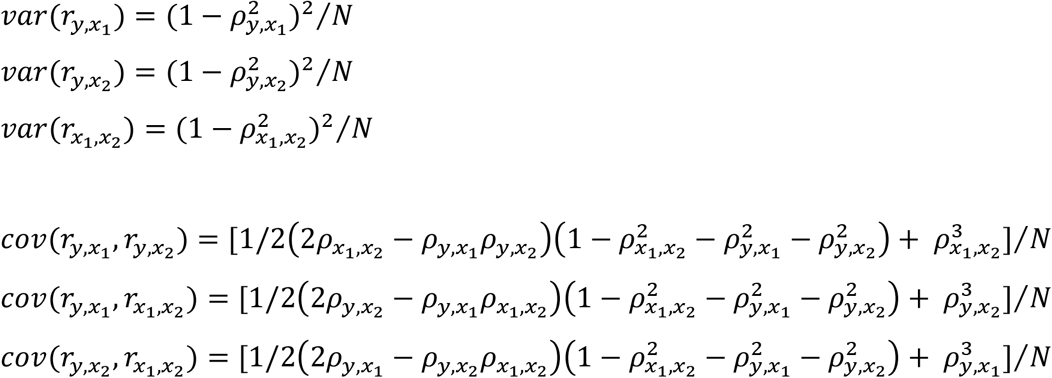

## Appendix B. r2redux manual

The ‘r2redux’ package can be used to derive test statistics for *R*^2^ values from polygenic risk score (PRS) models (variance and covariance of *R*^2^ values, p-value and 95% confidence intervals (CI)). For example, it can test if two sets of *R*^2^ values from two different PRS models are significantly different to each other whether the two sets of PRS are independent or dependent. Because *R*^2^ value is often regarded as the predictive ability of PRS, r2redux package can be useful to assess the performances of PRS methods or multiple sets of PRS based on different information sources. Furthermore, the package can derive the information matrix of 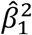and 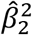 from a multiple regression (see olkin_beta1_2 or olkin_beta_info function in the manual), which is a basis of a novel PRS-based genomic partitioning method (see r2_enrich or r2_enrich_beta function in the manual). It is recommended that the target sample size in the PGS study should be more than 2,000 for quantitative traits (Supplementary Figure 27) and more than 5,000 for binary responses or case-control studies (Supplementary Figures 28 and 29). The p-value generated from r2redux is a two-tail test. Depending on hypothesis, one-tail p-value can be obtained as the two-tail p-value divided by 2.

### Installation

To use r2redux:

- `install.packages (“devtools”)`
- `library (devtools)`
- `devtools::install_github(“mommy003/r2redux_version4”) or`
- `install.packages (“r2redux”) [to be added in the CRAN soon]`
- `library (r2redux)`

### Quick start

We illustrate the usage of r2redux using multiple sets of PRS estimated based on GWAS summary statistics from UK Biobank or Biobank Japan (reference datasets). In a target dataset, the phenotypes of target samples (y) can be predicted with PRS (a PRS model, e.g. *y* = *PRS* + *e*, where y and PRS are column-standardised ^15^. Note that the target individuals should be independent from reference individuals. We can test the significant differences of the predictive ability (*R*^2^) between a pair of PRS (see r2_diff function and example in the manual).

### Data preparation

#### a. Statistical testing of significant difference between *R*^2^ values for p-value thresholds

r2redux requires only phenotype and estimated PRS (from PLINK or any other software). Note that any missing value in the phenotypes and PRS tested in the model should be removed. If we want to test the significant difference of *R*^2^ values for p-value thresholds, r2_diff function can be used with an input file that includes the following fields (also see test_ukbb_thresholds_scaled in the example directory form github (https://github.com/mommy003/r2redux_version4) and r2_diff function in the manual).

- Phenotype (y)
- PRS for p value 1 (*x*_1_)
- PRS for p value 0.5 (*x*_2_)
- PRS for p value 0.4 (*x*_3_)
- PRS for p value 0.3 (*x*_4_)
- PRS for p value 0.2 (*x*_5_)
- PRS for p value 0.1 (*x*_6_)
- PRS for p value 0.05 (*x*_7_)
- PRS for p value 0.01 (*x*_8_)
- PRS for p value 0.001 (*x*_9_)
- PRS for p value 0.0001 (*x*_10_)

To get the test statistics for the difference between *R*^2^(y~x[,v1]) and *R*^2^(y~x[,v2]). (here we define 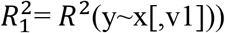 and 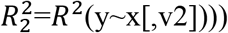

~~~
dat=read.table(“test_ukbb_thresholds_scaled”) (see example files)
nv=length(dat$V1)
v1=c(1)
v2=c(2)
output=r2_diff(dat,v1,v2,nv)
r2redux output
output$var1 (variance of 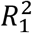)
0.0001437583
output$var2 (variance of 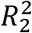)
0.0001452828
output$var_diff (variance of difference between 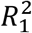 and 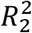)
5.678517e-07
output$r2_based_p (p-value for significant difference between 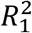 and 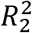)
0.5514562
output$mean_diff (differences between 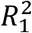 and 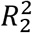)
-0.0004488044
output$upper_diff (upper limit of 95% CI for the difference)
0.001028172
output$lower_diff (lower limit of 95% CI for the difference)
-0.001925781
~~~

#### b. PRS-based genomic enrichment analysis

If we want to perform some enrichment analysis (e.g., regulatory vs non_regulatory) in the PRS context to test significantly different from the expectation (4%= # SNPs in the regulatory / total # SNPs). We simultaneously fit two sets of PRS from regulatory and non-regulatory to get 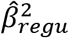 and 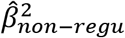, using a multiple regression, and assess if the ratio, 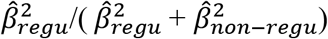 and/or 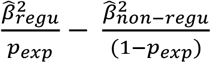, are significantly different from the expectation. To test this, we need to prepare input file for r2redux that includes the following fields (e.g. test_ukbb_enrichment_choles in example directory and r2_enrich_beta function in the manual).

- Phenotype (y)
- PRS for regulatory region (*x*_1_)
- PRS for non-regulatory region (*x*_2_)

To get the test statistic for the ratio which is significantly different from the expectation. 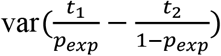, where 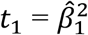 and 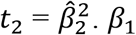 and *β*_2_ are regression coefficients from a multiple regression model, i.e. *y* = *x*_1_.*β*_1_ + *x*_2_.*β*_2_ + *e*, where *y*, *x*_1_ and *x*_2_ are column standardised.

~~~
dat=read.table(“test_ukbb_enrichment_choles”) (see example file)
nv=length(dat$V1)
v1=c(1)
v2=c(2)
expected_ratio=0.04
output=r2_enrich_beta(dat,v1,v2,nv,expected_ratio)
output
r2redux output
output$beta1_sq (*t*_1_)
0.01118301
output$beta2_sq (*t*_2_)
0.004980285
output$var1 (variance of *t*_1_)
7.072931e-05
output$var2 (variance of *t*_2_)
3.161929e-05
output$var1_2 (variance of difference between *t*_1_ and *t*_2_)
0.000162113
output$cov (covariance between *t*_1_ and *t*_2_)
-2.988221e-05
output$enrich_p2 (p-value for testing the difference between 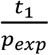 and 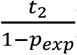)
0.1997805
output$mean_diff (difference between 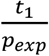 and 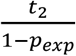)
0.2743874
output$var diff (variance of difference, 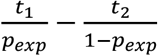)
0.04579649
output$upper_diff (upper limit of 95% CI for the mean difference)
0.6938296
output$lower_diff (lower limit of 95% CI for the mean difference)
-0.1450549
~~~

The r2redux manual and their example files can be downloaded from https://github.com/mommy003/r2redux_version4

