## Supplementary information for "Significance tests for *R*^2^ of out-of-sample prediction using polygenic scores"

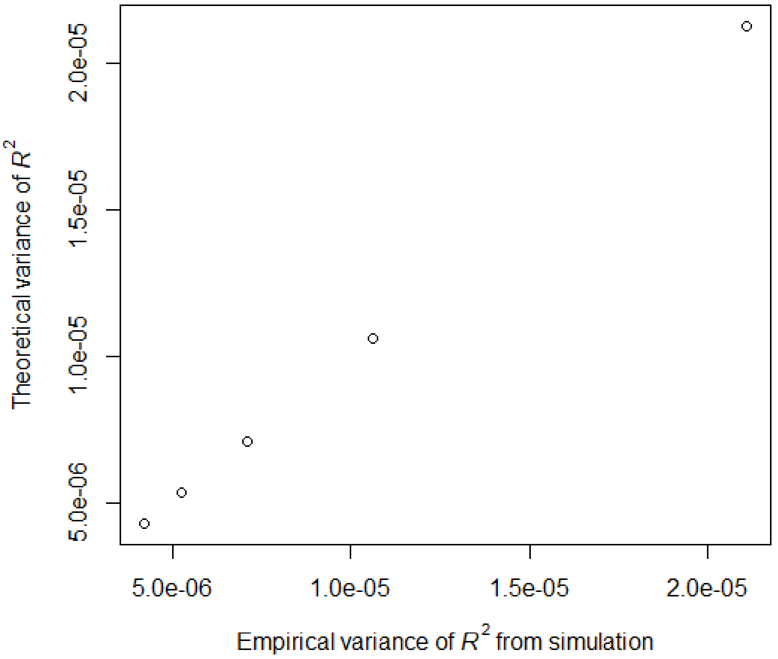


**Supplementary Figure 1: A near-perfect correlation between the theoretical and empirical variances of $R^2$ ($r^2_{y,x_1}$) estimated from 10,000 simulated replicates when varying sample size.** Simulations of $y$, $x_1$ and $x_2$ were based on a correlation structure $\begin{bmatrix} 1 & r_{y,x_1} & r_{y,x_2} \\ r_{y,x_1} & 1 & r_{x_1,x_2} \\ r_{y,x_2} & r_{x_1,x_2} & 1 \end{bmatrix} = \begin{bmatrix} 1 & 0.246 & 0.139 \\ 0.246 & 1 & 0.315 \\ 0.139 & 0.315 & 1 \end{bmatrix}$ and $R^2$ ($r^2_{y,x_1}$) was obtained from a model $y = x_1 + e$ in each replicate. The empirical variance of $R^2$ over 10,000 replicates was estimated. The theoretical variance of $R^2$ was obtained from eq. (5). Each data point in the diagonal represents the variance of $R^2$ with a sample size of 10000, 20000, 30000, 40000 and 50000.

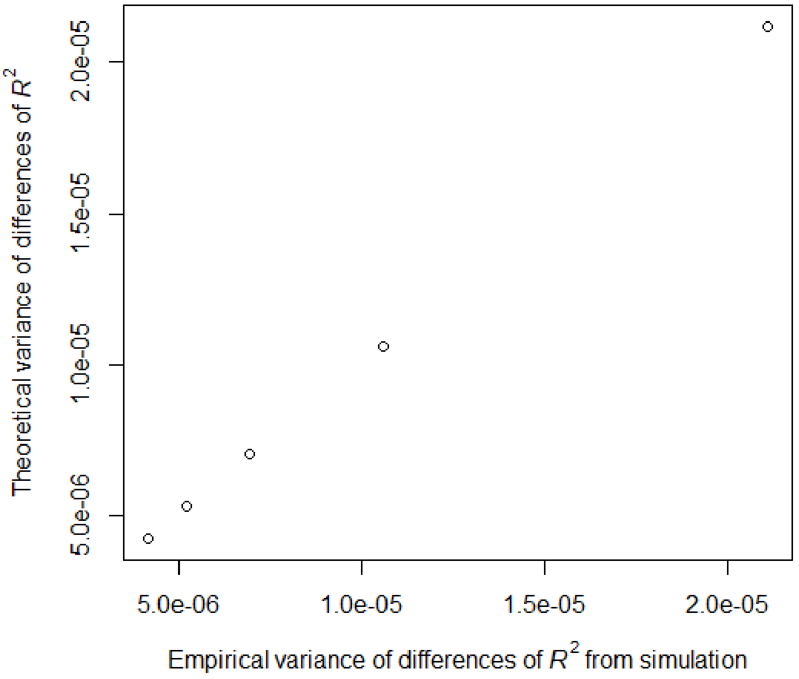


**Supplementary Figure 2: A near-perfect correlation between the theoretical and empirical variances of $R^2$ difference ($r^2_{y,x_1} - r^2_{y,x_2}$) estimated from 10,000 simulated replicates when varying sample size.** Simulations of $y$, $x_1$ and $x_2$ were based on a correlation structure $\begin{bmatrix} 1 & r_{y,x_1} & r_{y,x_2} \\ r_{y,x_1} & 1 & r_{x_1,x_2} \\ r_{y,x_2} & r_{x_1,x_2} & 1 \end{bmatrix} = \begin{bmatrix} 1 & 0.246 & 0.139 \\ 0.246 & 1 & 0.315 \\ 0.139 & 0.315 & 1 \end{bmatrix}$, and $r^2_{y,x_1}$ and $r^2_{y,x_2}$ were obtained from models $y = x_1 + e$ and $y = x_2 + e$, respectively, to get their difference in each replicate. The empirical variance of $r^2_{y,x_1} - r^2_{y,x_2}$ over 10,000 replicates was estimated. The theoretical variance of $r^2_{y,x_1} - r^2_{y,x_2}$ was obtained from eq. (9). Each data point in the diagonal represents the variance of $r^2_{y,x_1} - r^2_{y,x_2}$ with a sample size of 10000, 20000, 30000, 40000 and 50000.

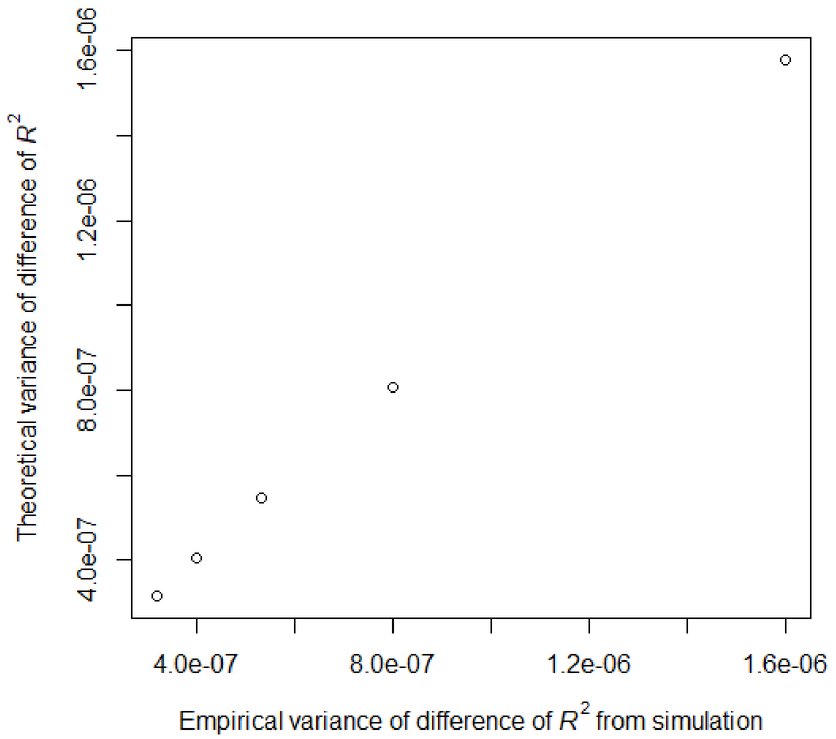


**Supplementary Figure 3: A near-perfect correlation between the theoretical and empirical variances of $R^2$ difference of $(r^2_{y,(x_1,x_2)} - r^2_{y,x_1})$ estimated from 10,000 simulated replicates when varying sample size.** Simulations of $y$, $x_1$ and $x_2$ were based on a correlation structure $\begin{bmatrix} 1 & r_{y,x_1} & r_{y,x_2} \\ r_{y,x_1} & 1 & r_{x_1,x_2} \\ r_{y,x_2} & r_{x_1,x_2} & 1 \end{bmatrix} = \begin{bmatrix} 1 & 0.246 & 0.139 \\ 0.246 & 1 & 0.315 \\ 0.139 & 0.315 & 1 \end{bmatrix}$, and $r^2_{y,(x_1,x_2)}$ and $r^2_{y,x_1}$ were obtained from models $y = x_1 + x_2 + e$ and $y = x_1 + e$, respectively, to get their difference in each replicate. The empirical variance of $r^2_{y,(x_1,x_2)} - r^2_{y,x_1}$ over 10,000 replicates was estimated. The theoretical variance of $r^2_{y,(x_1,x_2)} - r^2_{y,x_1}$ was obtained from eq. (11). Each data point in the diagonal represents the variance of $r^2_{y,(x_1,x_2)} - r^2_{y,x_1}$ with a sample size of 10000, 20000, 30000, 40000 and 50000.

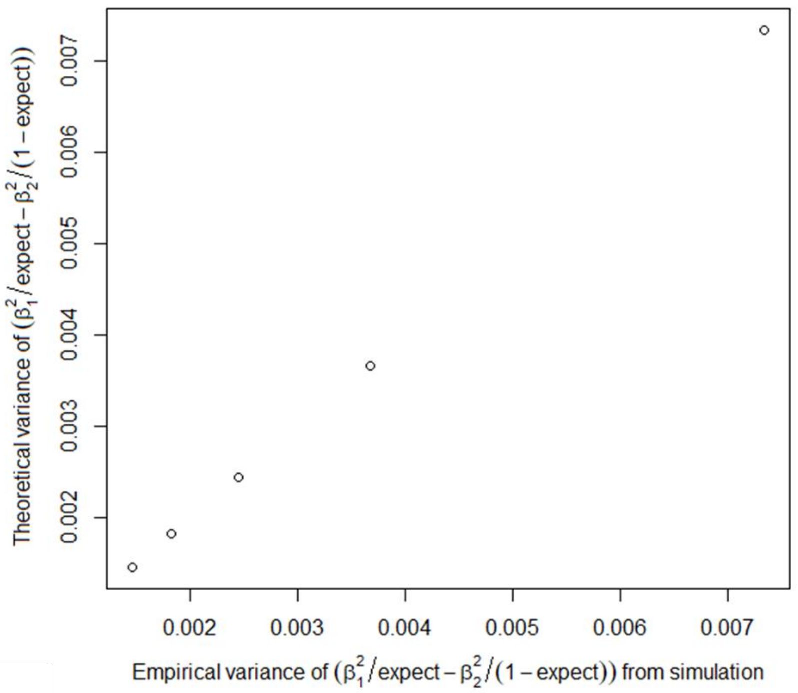


**Supplementary Figure 4: A near-perfect correlation between the theoretical and empirical variances of $\frac{\hat{\beta}_1^2}{p_{exp}} - \frac{\hat{\beta}_2^2}{(1-p_{exp})}$ estimated from 10,000 simulated replicates when varying sample size.**

Simulations of $y$, $x_1$ and $x_2$ were based on a correlation structure $\begin{bmatrix} 1 & r_{y,x_1} & r_{y,x_2} \\ r_{y,x_1} & 1 & r_{x_1,x_2} \\ r_{y,x_2} & r_{x_1,x_2} & 1 \end{bmatrix} = \begin{bmatrix} 1 & 0.176 & 0.148 \\ 0.176 & 1 & 0.610 \\ 0.148 & 0.610 & 1 \end{bmatrix}$, and $\hat{\beta}_1^2$ and $\hat{\beta}_2^2$ were obtained from a multiple regression model $y = x_1 + x_2 + e$ to get the difference of scaled squared regression coefficients in each replicate. It was assumed that the expectation is known ($p_{exp}$= 0.04 was used). The empirical variance of $\frac{\hat{\beta}_1^2}{p_{exp}} - \frac{\hat{\beta}_2^2}{(1-p_{exp})}$ over 10,000 replicates was estimated. The theoretical variance of $\frac{\hat{\beta}_1^2}{p_{exp}} - \frac{\hat{\beta}_2^2}{(1-p_{exp})}$ was obtained from eq. (17). Each data point in the diagonal represents the variance of $\frac{\hat{\beta}_1^2}{p_{exp}} - \frac{\hat{\beta}_2^2}{(1-p_{exp})}$ with a sample size of 10000, 20000, 30000, 40000 and 50000.

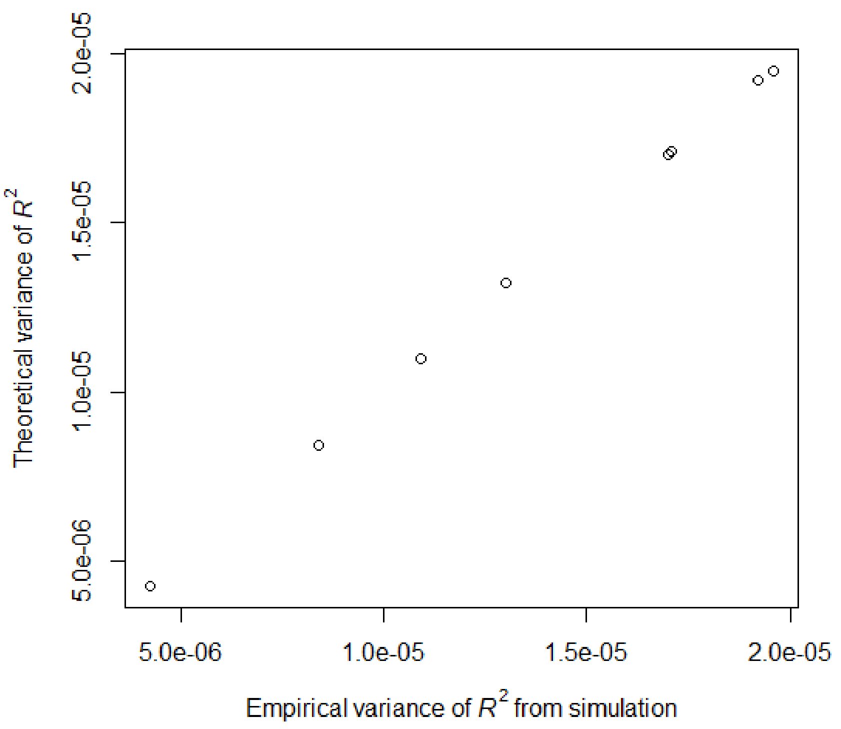


**Supplementary Figure 5: A near-perfect correlation between the theoretical and empirical variances of $R^2$ ($r^2_{y,x_1}$) estimated from 10,000 simulated replicates when varying $R^2$ value.** Simulations of $y$, $x_1$ and $x_2$ were based on a correlation structure $\begin{bmatrix} 1 & r_{y,x_1} & r_{y,x_2} \\ r_{y,x_1} & 1 & r_{x_1,x_2} \\ r_{y,x_2} & r_{x_1,x_2} & 1 \end{bmatrix} = \begin{bmatrix} 1 & various & 0.447 \\ various & 1 & 0.800 \\ 0.447 & 0.800 & 1 \end{bmatrix}$ and $R^2$ ($r^2_{y,x_1}$) was obtained from a model $y = x_1 + e$ in each replicate. The empirical variance of $R^2$ over 10,000 replicates was estimated. The theoretical variance of $R^2$ was obtained from eq. (5). A sample size of 30,000 was used. Each data point in the diagonal represents the variance of $R^2$ with $r^2_{y,x_1}$ = 0.1, 0.2, 0.3, 0.4, 0.5, 0.6, 0.7 and 0.8.

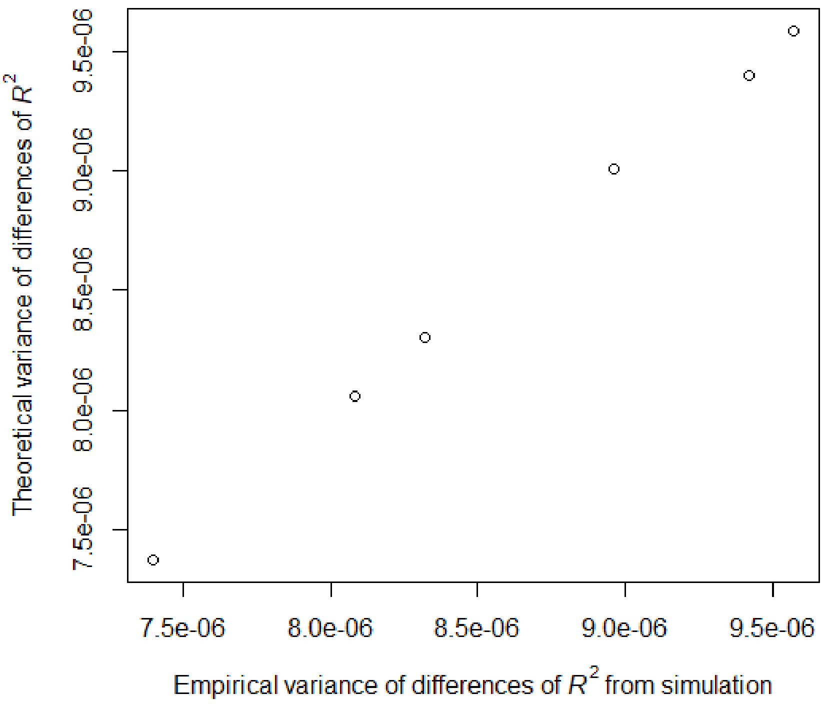


**Supplementary Figure 6: A near-perfect correlation between the theoretical and empirical variances of $R^2$ difference ($r^2_{y,x_1} - r^2_{y,x_2}$) estimated from 10,000 simulated replicates when varying $R^2$ difference.** Simulations of $y$, $x_1$ and $x_2$ were based on a correlation structure $\begin{bmatrix} 1 & r_{y,x_1} & r_{y,x_2} \\ r_{y,x_1} & 1 & r_{x_1,x_2} \\ r_{y,x_2} & r_{x_1,x_2} & 1 \end{bmatrix} = \begin{bmatrix} 1 & various & 0.447 \\ various & 1 & 0.800 \\ 0.447 & 0.800 & 1 \end{bmatrix}$, and $r^2_{y,x_1}$ and $r^2_{y,x_2}$ were obtained from models $y = x_1 + e$ and $y = x_2 + e$, respectively, to get their difference in each replicate. The empirical variance of $r^2_{y,x_1} - r^2_{y,x_2}$ over 10,000 replicates was estimated. The theoretical variance of $r^2_{y,x_1} - r^2_{y,x_2}$ was obtained from eq. (9). A sample size of 30,000 was used. Each data point in the diagonal represents the variance of $r^2_{y,x_1} - r^2_{y,x_2}$ with $r^2_{y,x_1} - r^2_{y,x_2}$ = 0, 0.1, 0.2, 0.3, 0.4 and 0.5.

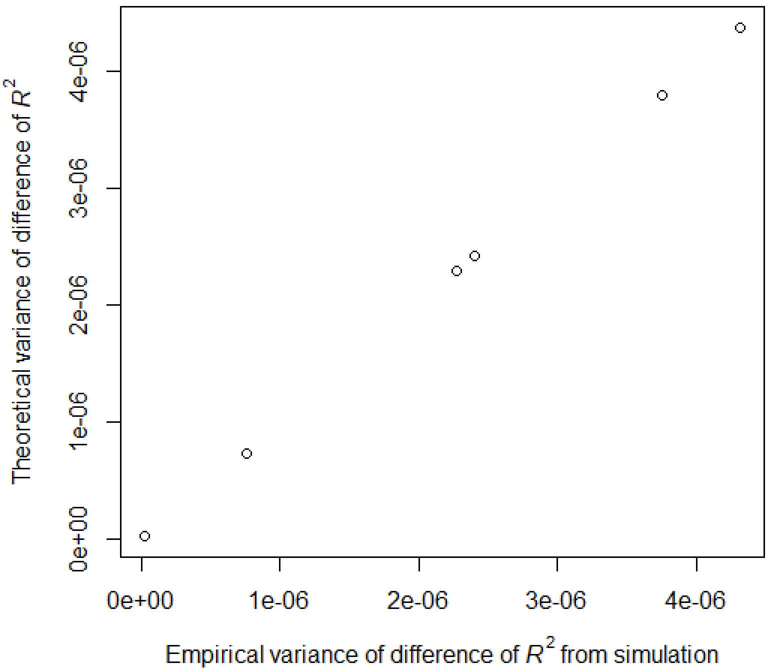


**Supplementary Figure 7: A near-perfect correlation between the theoretical and empirical variances of $R^2$ difference ($r^2_{y,(x_1,x_2)} - r^2_{y,x_1}$) estimated from 10,000 simulated replicates when varying $R^2$ difference.** Simulations of $y$, $x_1$ and $x_2$ were based on a correlation structure $\begin{bmatrix} 1 & r_{y,x_1} & r_{y,x_2} \\ r_{y,x_1} & 1 & r_{x_1,x_2} \\ r_{y,x_2} & r_{x_1,x_2} & 1 \end{bmatrix} = \begin{bmatrix} 1 & various & 0.447 \\ various & 1 & 0.800 \\ 0.447 & 0.800 & 1 \end{bmatrix}$, and $r^2_{y,(x_1,x_2)}$ and $r^2_{y,x_1}$ were obtained from models $y = x_1 + x_2 + e$ and $y = x_1 + e$, respectively, to get their difference in each replicate. The empirical variance of $r^2_{y,(x_1,x_2)} - r^2_{y,x_1}$ over 10,000 replicates was estimated. The theoretical variance of$r^2_{y,(x_1,x_2)} - r^2_{y,x_1}$ was obtained from eq. (11). A sample size of 30,000 was used. Each data point in the diagonal represents the variance of $r^2_{y,(x_1,x_2)} - r^2_{y,x_1}$ with $r^2_{y,(x_1,x_2)} - r^2_{y,x_1}$ = 0, 0.1, 0.2, 0.3, 0.4 and 0.5.

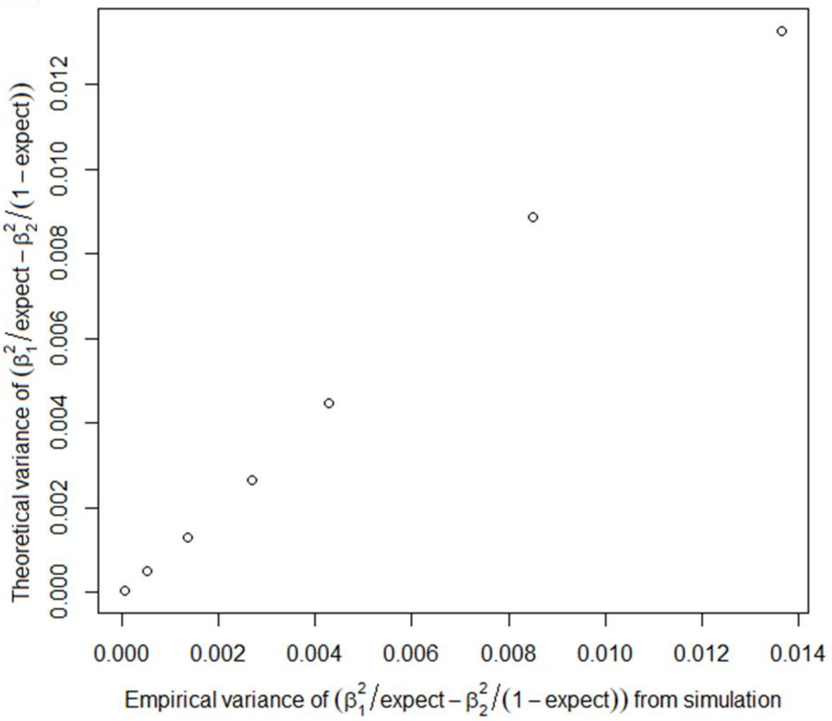


**Supplementary Figure 8. A near-perfect correlation between the theoretical and empirical variances of $\frac{\hat{\beta}_1^2}{p_{exp}} - \frac{\hat{\beta}_2^2}{(1-p_{exp})}$ estimated from 10,000 simulated replicates when varying correlation structure.** Simulations of $y$, $x_1$ and $x_2$ were based on a correlation structure $\begin{bmatrix} 1 & r_{y,x_1} & r_{y,x_2} \\ r_{y,x_1} & 1 & r_{x_1,x_2} \\ r_{y,x_2} & r_{x_1,x_2} & 1 \end{bmatrix} = \begin{bmatrix} 1 & various & 0.148 \\ various & 1 & 0.610 \\ 0.148 & 0.610 & 1 \end{bmatrix}$, and $\hat{\beta}_1^2$ and $\hat{\beta}_2^2$ were obtained from a multiple regression model $y = x_1 + x_2 + e$ to get the difference of scaled squared regression coefficients in each replicate. It was assumed that the expectation is known ($p_{exp}$ = 0.04 was used). The empirical variance of $\frac{\hat{\beta}_1^2}{p_{exp}} - \frac{\hat{\beta}_2^2}{(1-p_{exp})}$ over 10,000 replicates was estimated. The theoretical variance of $\frac{\hat{\beta}_1^2}{p_{exp}} - \frac{\hat{\beta}_2^2}{(1-p_{exp})}$ was obtained from eq. (17). A sample size of 30,000 was used. Each data point in the diagonal represents the variance of $\frac{\hat{\beta}_1^2}{p_{exp}} - \frac{\hat{\beta}_2^2}{(1-p_{exp})}$ with $r_{y,x_1}$ = 0.05, 0.10, 0.15, 0.20, 0.25 and 0.30 (resulting in $\frac{\hat{\beta}_1^2}{p_{exp}} - \frac{\hat{\beta}_2^2}{(1-p_{exp})}$ = 0.067, 0.013, 0.220, 0.767, 1.619 and 2.786).

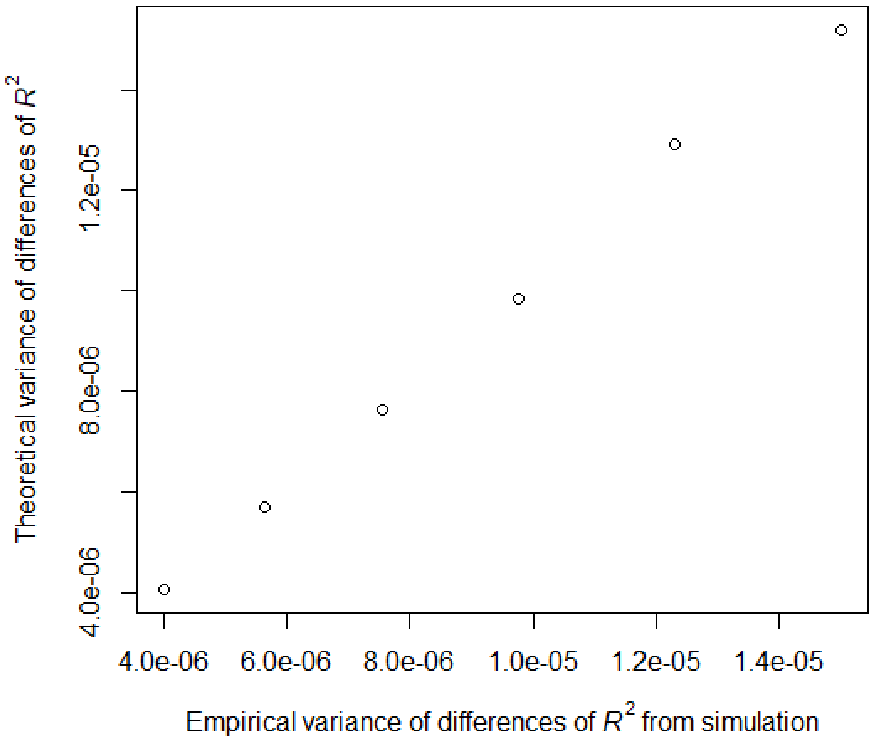


**Supplementary Figure 9: A near-perfect correlation between the theoretical and empirical variances of $R^2$ difference ($r^2_{y_1,x_1} - r^2_{y_2,x_2}$) estimated from 10,000 simulated replicates when using two sets of independent PGS (e.g., Male vs female PGS).** Simulations of $y_1$ and $x_1$ were based on a correlation structure $\begin{bmatrix} 1 & r_{y_1,x_1} \\ r_{y_1,x_1} & 1 \end{bmatrix} = \begin{bmatrix} 1 & various \\ various & 1 \end{bmatrix}$, simulations of $y_2$ and $x_2$ were based on a correlation structure $\begin{bmatrix} 1 & r_{y_2,x_2} \\ r_{y_2,x_2} & 1 \end{bmatrix} = \begin{bmatrix} 1 & 0.447 \\ 0.447 & 1 \end{bmatrix}$ and $r^2_{y_1,x_1}$ and $r^2_{y_2,x_2}$ were obtained from models $y_1 = x_1 + e$ and $y_2 = x_2 + e$, respectively, to get their difference in each replicate. The empirical variance of $r^2_{y1,x_1} - r^2_{y2,x_2}$ over 10,000 replicates was estimated. The theoretical variance of $r^2_{y_1,x_1} - r^2_{y_2,x_2}$ was obtained from eq. (14). A sample size of 15,000 and 17,000 were used for 1st and 2nd PGS, respectively. Each data point in the diagonal represents the variance of $r^2_{y_1,x_1} - r^2_{y_2,x_2}$ with $r^2_{y_1,x_1} - r^2_{y_2,x_2}$ = 0, 0.02, 0.04, 0.06, 0.08 and 0.10.

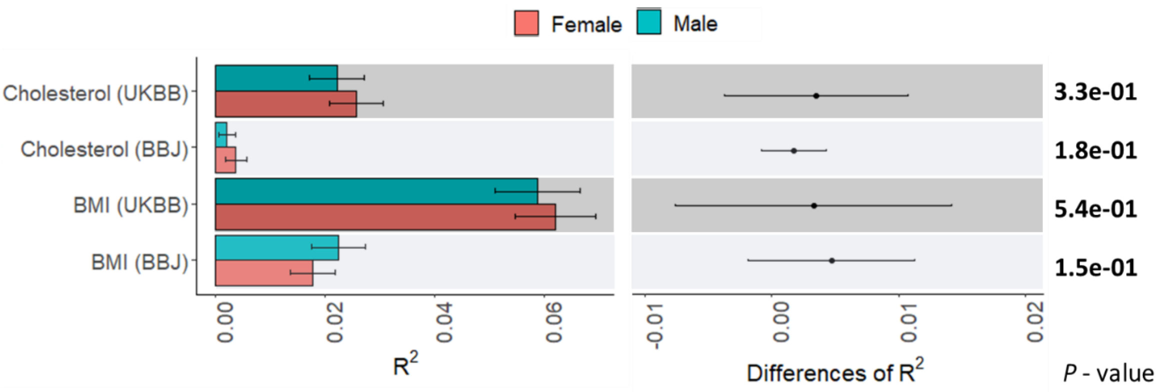


**Supplementary Figure 10: The predictive ability of ($R^2$) of male and female, when predicting European male and female separately using UKBB and BBJ discovery samples.**
**Left panel:** The main bars represent $R^2$ values and error bars correspond 95% confidence intervals.
**Right panel:** Dot points represent the differences of $R^2$ values between male and female PGS models, and error bars indicate 95% confidence intervals of the difference.
95% confidence interval for the differences of $R^2$ between two independent sets of PGS (male and female) was estimated from eq. (15).

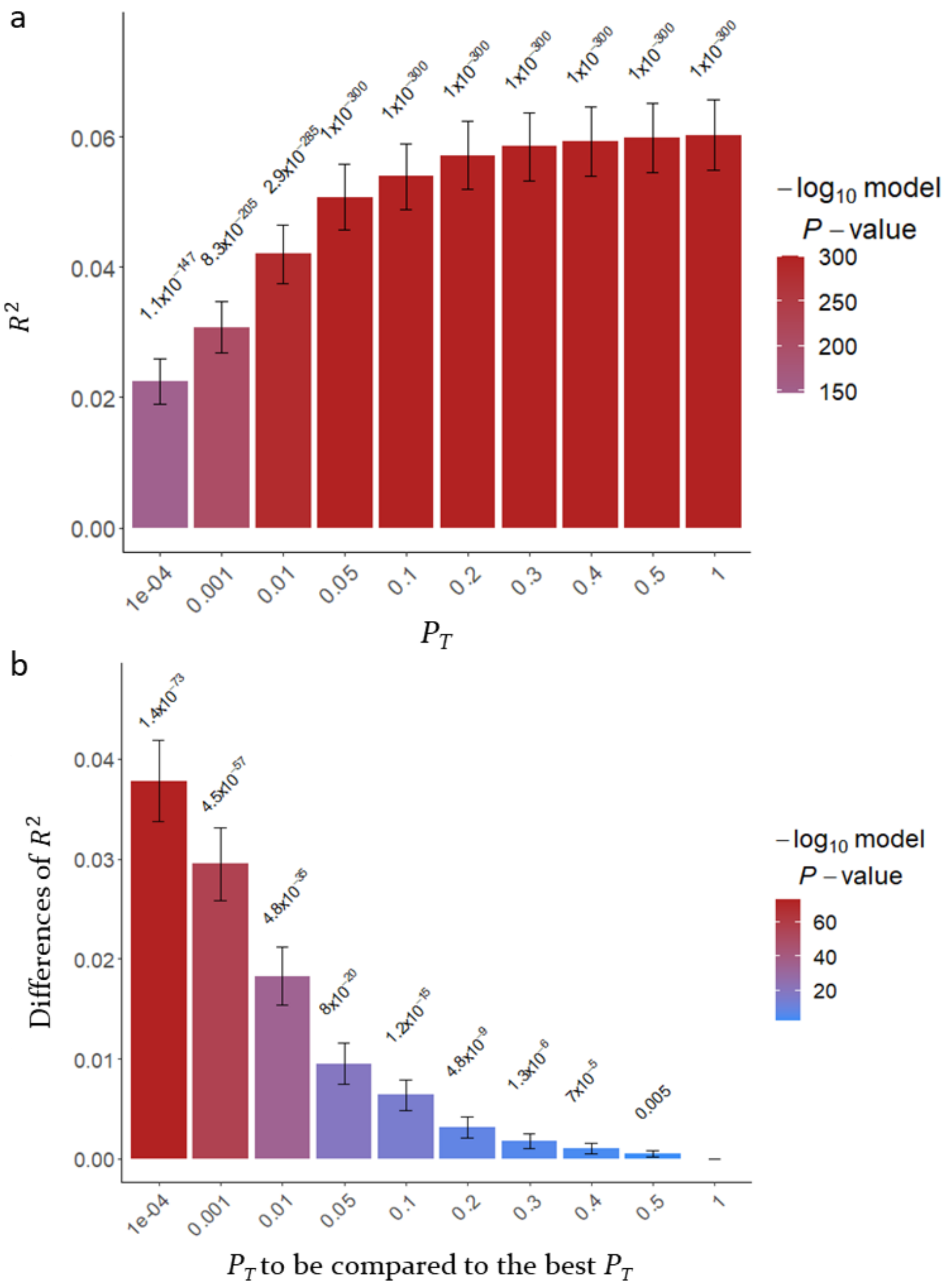


**Supplementary Figure 11: The predictive ability ($R^2$) of PGS estimated based on SNPs below the $p_T$ when predicting BMI in 28,880 European samples using UKBB discovery samples (GWAS summary statistics).**

a) The main bars represent $R^2$ values and error bars correspond 95% confidence intervals. The values above 95% CIs are p-values indicating that $R^2$ values are not different from zero.

b) The main bars represent the difference of $R^2$ values between the corresponding threshold and the best-performed threshold and error bars indicate 95% confidence intervals. The values above 95% CIs are p-values indicating the significances of differences between the pairs of $R^2$ values.

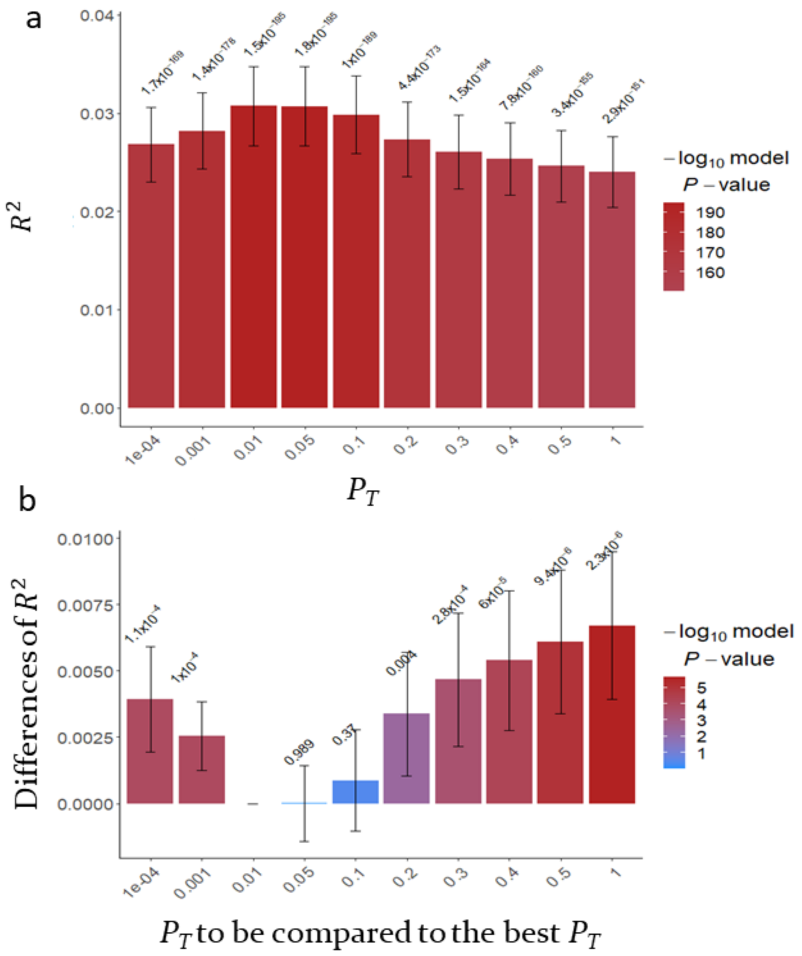


**Supplementary Figure 12: The predictive ability ($R^2$) of PGS estimated based on SNPs below the $p_T$ when predicting cholesterol in 28,880 European samples using UKBB discovery samples (GWAS summary statistics).**

a) The main bars represent $R^2$ values and error bars correspond 95% confidence intervals. The values above 95% CIs are p-values indicating that $R^2$ values are not different from zero.

b) The main bars represent the difference of $R^2$ values between the corresponding threshold and the best-performed threshold and error bars indicate 95% confidence intervals. The values above 95% CIs are p-values indicating the significances of differences between the pairs of $R^2$ values.

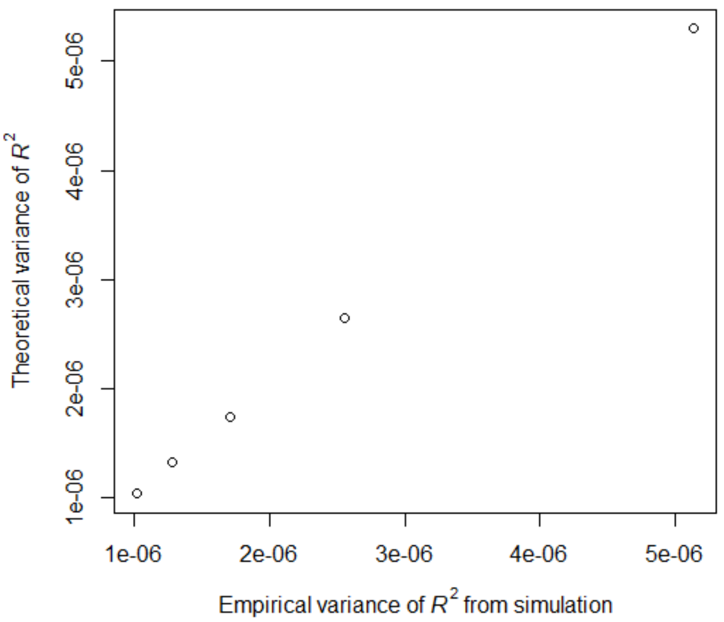


**Supplementary Figure 13: A near-perfect correlation between the theoretical and empirical variances of $R^2$ ($r^2_{y,x_1}$) estimated from 10,000 simulated replicates of binary responses assuming 5% disease prevalence when varying sample size.** Simulations of $y$, $x_1$ and $x_2$ were based on a correlation structure $\begin{bmatrix} 1 & r_{y,x_1} & r_{y,x_2} \\ r_{y,x_1} & 1 & r_{x_1,x_2} \\ r_{y,x_2} & r_{x_1,x_2} & 1 \end{bmatrix} = \begin{bmatrix} 1 & 0.246 & 0.139 \\ 0.246 & 1 & 0.315 \\ 0.139 & 0.315 & 1 \end{bmatrix}$ and $R^2$ ($r^2_{y,x_1}$) was obtained from a model $y = x_1 + e$ in each replicate. The empirical variance of $R^2$ over 10,000 replicates was estimated. The theoretical variance of $R^2$ was obtained from eq. (5). Each data point in the diagonal represents the variance of $R^2$ with a sample size of 10000, 20000, 30000, 40000 and 50000.

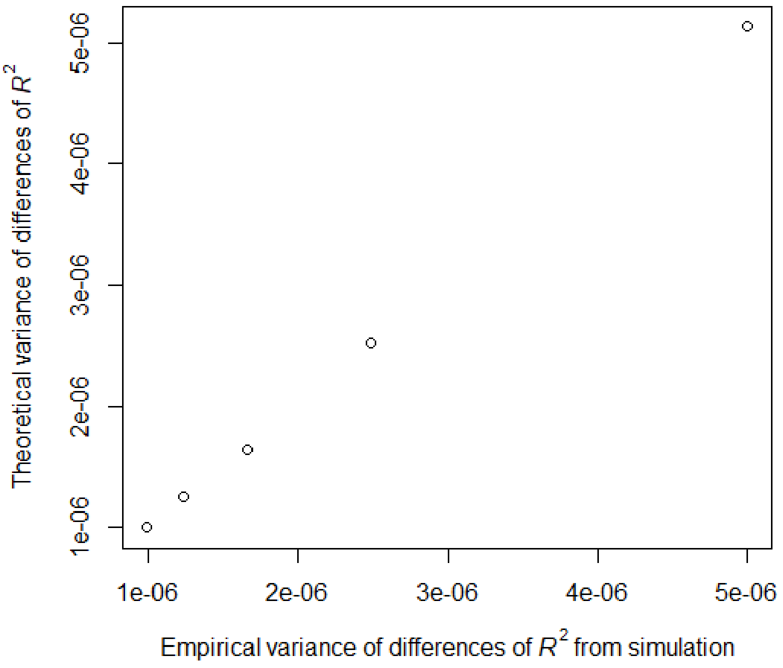


**Supplementary Figure 14: A near-perfect correlation between the theoretical and empirical variances of $R^2$ difference ($r^2_{y,x_1} - r^2_{y,x_2}$) estimated from 10,000 simulated replicates of binary responses assuming 5% disease prevalence when varying sample size.** Simulations of $y$, $x_1$ and $x_2$ were based on a correlation structure $\begin{bmatrix} 1 & r_{y,x_1} & r_{y,x_2} \\ r_{y,x_1} & 1 & r_{x_1,x_2} \\ r_{y,x_2} & r_{x_1,x_2} & 1 \end{bmatrix} = \begin{bmatrix} 1 & 0.246 & 0.139 \\ 0.246 & 1 & 0.315 \\ 0.139 & 0.315 & 1 \end{bmatrix}$, and $r^2_{y,x_1}$ and $r^2_{y,x_2}$ were obtained from models $y = x_1 + e$ and $y = x_2 + e$, respectively, to get their difference in each replicate. The empirical variance of $r^2_{y,x_1} - r^2_{y,x_2}$ over 10,000 replicates was estimated. The theoretical variance of $r^2_{y,x_1} - r^2_{y,x_2}$ was obtained from eq. (9). Each data point in the diagonal represents the variance of $r^2_{y,x_1} - r^2_{y,x_2}$ with a sample size of 10000, 20000, 30000, 40000 and 50000.

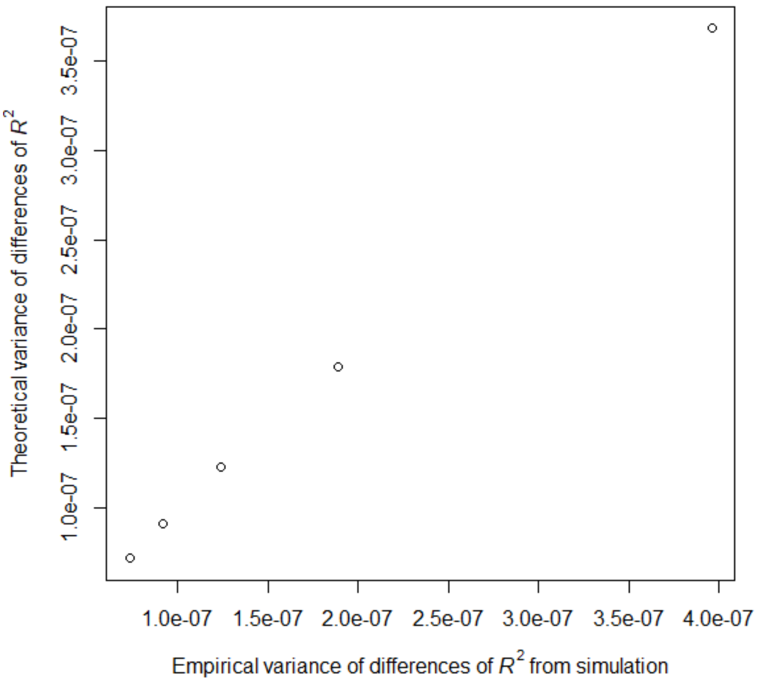


**Supplementary Figure 15: A near-perfect correlation between the theoretical and empirical variances of $R^2$ difference ($r^2_{y,(x_1,x_2)} - r^2_{y,x_1}$) estimated from 10,000 simulated replicates of binary responses assuming 5% disease prevalence when varying sample size.** Simulations of $y$, $x_1$ and $x_2$ were based on a correlation structure $\begin{bmatrix} 1 & r_{y,x_1} & r_{y,x_2} \\ r_{y,x_1} & 1 & r_{x_1,x_2} \\ r_{y,x_2} & r_{x_1,x_2} & 1 \end{bmatrix} = \begin{bmatrix} 1 & 0.246 & 0.139 \\ 0.246 & 1 & 0.315 \\ 0.139 & 0.315 & 1 \end{bmatrix}$, and $r^2_{y,(x_1,x_2)}$ and $r^2_{y,x_1}$ were obtained from models $y = x_1 + x_2 + e$ and $y = x_1 + e$, respectively, to get their difference in each replicate. The empirical variance of $r^2_{y,(x_1,x_2)} - r^2_{y,x_1}$ over 10,000 replicates was estimated. The theoretical variance of $r^2_{y,(x_1,x_2)} - r^2_{y,x_1}$ was obtained from eq. (11). Each data point in the diagonal represents the variance of $r^2_{y,(x_1,x_2)} - r^2_{y,x_1}$ with a sample size of 10000, 20000, 30000, 40000 and 50000.

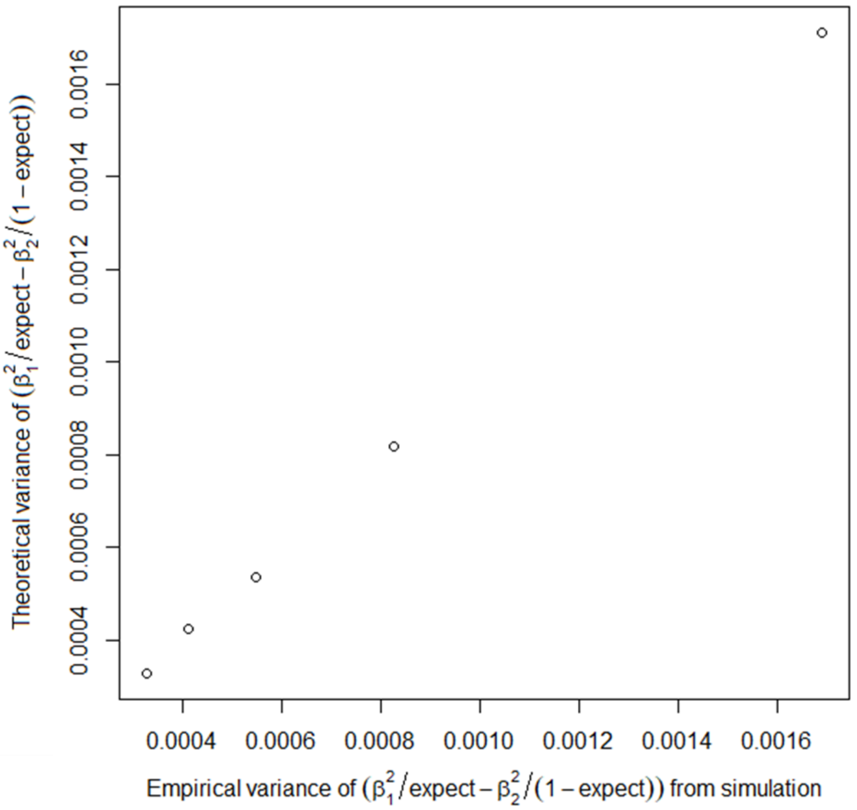


**Supplementary Figure 16: A near-perfect correlation between the theoretical and empirical variances of $\frac{\hat{\beta}_1^2}{p_{exp}} - \frac{\hat{\beta}_2^2}{(1-p_{exp})}$ estimated from 10,000 simulated replicates of binary responses assuming 5% disease prevalence when varying sample size.** Simulations of $y$, $x_1$ and $x_2$ were based on a correlation structure $\begin{bmatrix} 1 & r_{y,x_1} & r_{y,x_2} \\ r_{y,x_1} & 1 & r_{x_1,x_2} \\ r_{y,x_2} & r_{x_1,x_2} & 1 \end{bmatrix} = \begin{bmatrix} 1 & 0.176 & 0.148 \\ 0.176 & 1 & 0.610 \\ 0.148 & 0.610 & 1 \end{bmatrix}$, and $\hat{\beta}_1^2$ and $\hat{\beta}_2^2$ were obtained from a multiple regression model $y = x_1 + x_2 + e$ to get the difference of scaled squared regression coefficients in each replicate. It was assumed that the expectation is known ($p_{exp}$ = 0.04 was used). The empirical variance of $\frac{\hat{\beta}_1^2}{p_{exp}} - \frac{\hat{\beta}_2^2}{(1-p_{exp})}$ over 10,000 replicates was estimated. The theoretical variance of $\frac{\hat{\beta}_1^2}{p_{exp}} - \frac{\hat{\beta}_2^2}{(1-p_{exp})}$ was obtained from eq. (17). Each data point in the diagonal represents the variance of $\frac{\hat{\beta}_1^2}{p_{exp}} - \frac{\hat{\beta}_2^2}{(1-p_{exp})}$ with a sample size of 10000, 20000, 30000, 40000 and 50000.

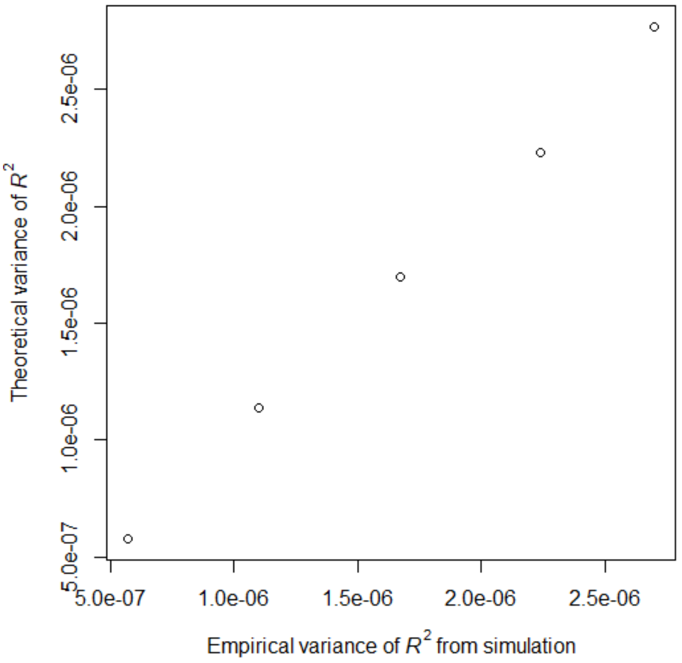


**Supplementary Figure 17: A near-perfect correlation between the theoretical and empirical variances of $R^2$ ($r^2_{y,x_1}$) estimated from 10,000 simulated replicates of binary responses assuming 5% disease prevalence when varying $R^2$ value.** Simulations of $y$, $x_1$ and $x_2$ were based on a correlation structure $\begin{bmatrix} 1 & r_{y,x_1} & r_{y,x_2} \\ r_{y,x_1} & 1 & r_{x_1,x_2} \\ r_{y,x_2} & r_{x_1,x_2} & 1 \end{bmatrix} = \begin{bmatrix} 1 & various & 0.141 \\ various & 1 & 0.800 \\ 0.141 & 0.8 & 1 \end{bmatrix}$ and $R^2(r^2_{y,x_1})$ was obtained from a model $y = x_1 + e$ in each replicate. The empirical variance of $R^2$ over 10,000 replicates was estimated. The theoretical variance of $R^2$ was obtained from eq. (5). A sample size of 30,000 was used. Each data point in the diagonal represents the variance of $R^2$ with $r^2_{y,x_1}$ = 0.02, 0.04, 0.06, 0.08, and 0.10.

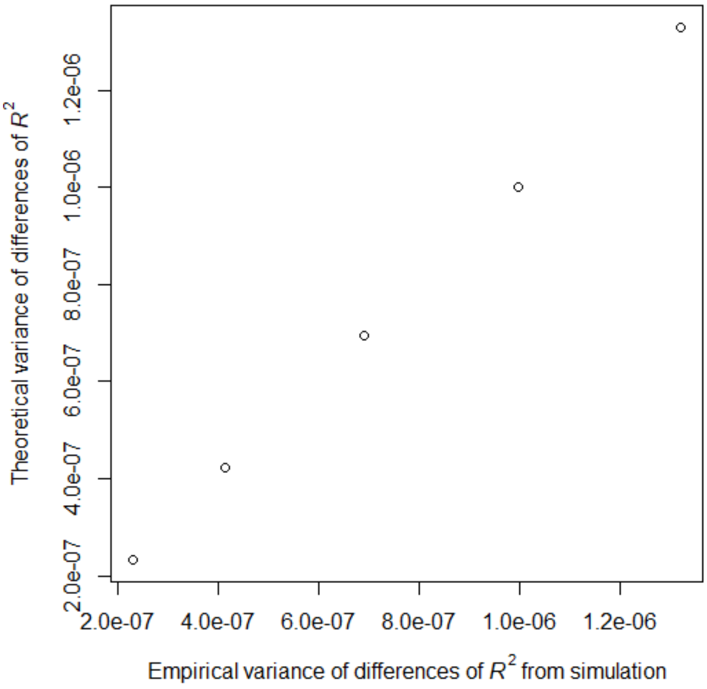


**Supplementary Figure 18: A near-perfect correlation between the theoretical and empirical variances of $R^2$ difference ($r^2_{y,x_1} - r^2_{y,x_2}$) estimated from 10,000 simulated replicates of binary responses assuming 5% disease prevalence when varying $R^2$ difference.** Simulations of $y$, $x_1$ and $x_2$ were based on a correlation structure $\begin{bmatrix} 1 & r_{y,x_1} & r_{y,x_2} \\ r_{y,x_1} & 1 & r_{x_1,x_2} \\ r_{y,x_2} & r_{x_1,x_2} & 1 \end{bmatrix} = \begin{bmatrix} 1 & various & 0.141 \\ various & 1 & 0.800 \\ 0.141 & 0.8 & 1 \end{bmatrix}$, and $r^2_{y,x_1}$ and $r^2_{y,x_2}$ were obtained from models $y = x_1 + e$ and $y = x_2 + e$, respectively, to get their difference in each replicate. The empirical variance of $r^2_{y,x_1} - r^2_{y,x_2}$ over 10,000 replicates was estimated. The theoretical variance of $r^2_{y,x_1} - r^2_{y,x_2}$ was obtained from eq. (9). A sample size of 30,000 was used. Each data point in the diagonal represents the variance of $r^2_{y,x_1} - r^2_{y,x_2}$ with $r^2_{y,x_1} - r^2_{y,x_2}$ = 0, 0.02, 0.04, 0.0.06, and0.08.

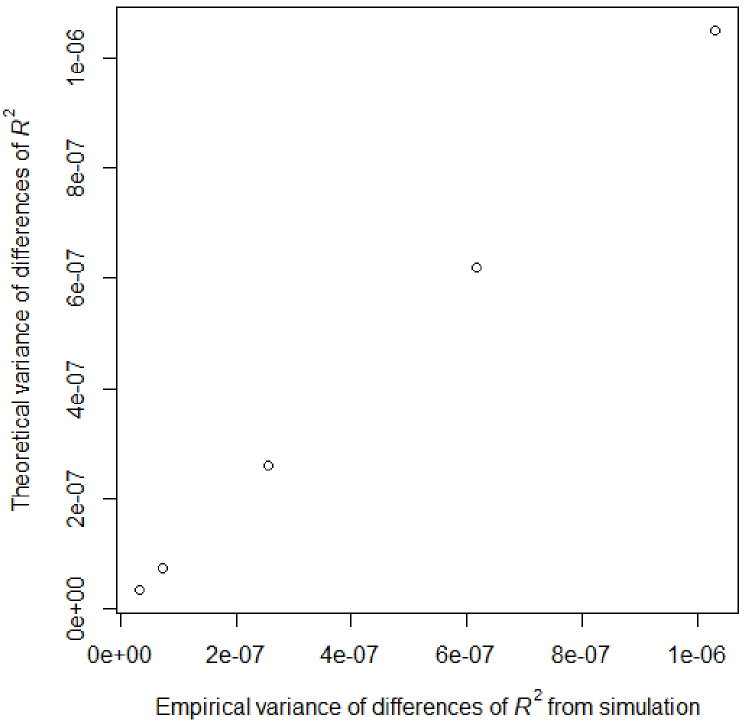


**Supplementary Figure 19: A near-perfect correlation between the theoretical and empirical variances of $R^2$ difference ($r^2_{y,(x_1,x_2)} - r^2_{y,x_1}$) estimated from 10,000 simulated replicates of binary responses assuming 5% disease prevalence when varying $R^2$ difference.** Simulations of $y$, $x_1$ and $x_2$ were based on a correlation structure $\begin{bmatrix} 1 & r_{y,x_1} & r_{y,x_2} \\ r_{y,x_1} & 1 & r_{x_1,x_2} \\ r_{y,x_2} & r_{x_1,x_2} & 1 \end{bmatrix} = \begin{bmatrix} 1 & various & 0.141 \\ various & 1 & 0.800 \\ 0.141 & 0.8 & 1 \end{bmatrix}$, and $r^2_{y,(x_1,x_2)}$ and $r^2_{y,x_1}$ were obtained from models $y = x_1 + x_2 + e$ and $y = x_1 + e$, respectively, to get their difference in each replicate. The empirical variance of $r^2_{y,(x_1,x_2)} - r^2_{y,x_1}$ over 10,000 replicates was estimated. The theoretical variance of$r^2_{y,(x_1,x_2)} - r^2_{y,x_1}$ was obtained from eq. (11). A sample size of 30,000 was used. Each data point in the diagonal represents the variance of $r^2_{y,(x_1,x_2)} - r^2_{y,x_1}$ with $r^2_{y,(x_1,x_2)} - r^2_{y,x_1}$ = 0, 0.02, 0.04, 0.0.06, and 0.08.

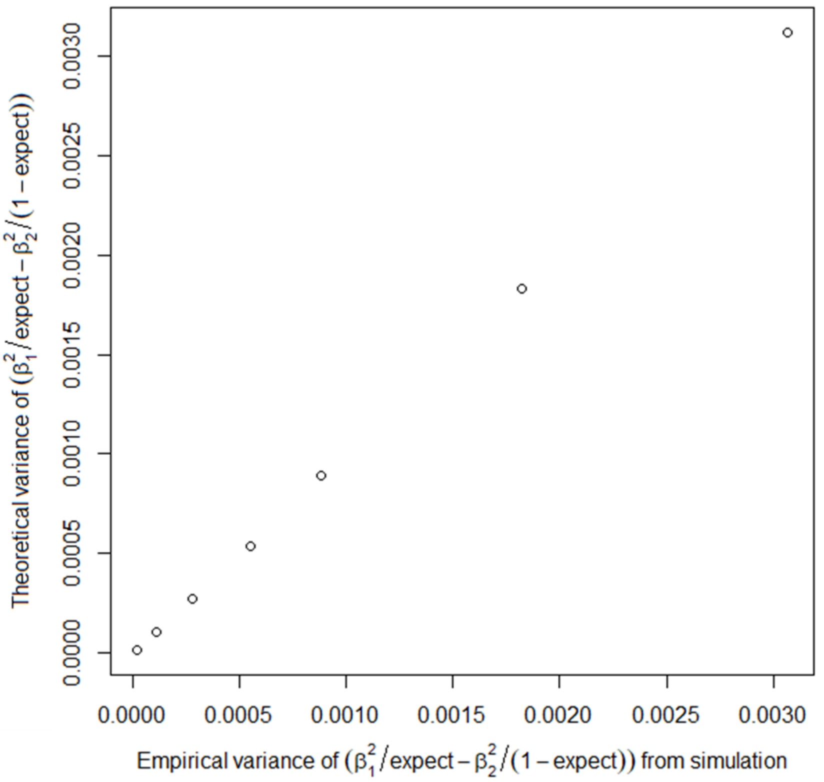


**Supplementary Figure 20: A near-perfect correlation between the theoretical and empirical variances of $\frac{\hat{\beta}_1^2}{p_{exp}} - \frac{\hat{\beta}_2^2}{(1-p_{exp})}$ estimated from 10,000 simulated replicates of binary responses assuming 5% disease prevalence when varying correlation structure.** Simulations of $y$, $x_1$ and $x_2$ were based on a correlation structure $\begin{bmatrix} 1 & r_{y,x_1} & r_{y,x_2} \\ r_{y,x_1} & 1 & r_{x_1,x_2} \\ r_{y,x_2} & r_{x_1,x_2} & 1 \end{bmatrix} = \begin{bmatrix} 1 & various & 0.148 \\ various & 1 & 0.610 \\ 0.148 & 0.610 & 1 \end{bmatrix}$, and $\hat{\beta}_1^2$ and $\hat{\beta}_2^2$ were obtained from a multiple regression model $y = x_1 + x_2 + e$ to get the difference of scaled squared regression coefficients in each replicate. It was assumed that the expectation is known ($p_{exp}$ = 0.04 was used). The empirical variance of $\frac{\hat{\beta}_1^2}{p_{exp}} - \frac{\hat{\beta}_2^2}{(1-p_{exp})}$ over 10,000 replicates was estimated. The theoretical variance of $\frac{\hat{\beta}_1^2}{p_{exp}} - \frac{\hat{\beta}_2^2}{(1-p_{exp})}$ was obtained from eq. (17). A sample size of 30,000 was used. Each data point in the diagonal represents the **variance of $\frac{\hat{\beta}_1^2}{p_{exp}} - \frac{\hat{\beta}_2^2}{(1-p_{exp})}$ with $r_{y,x_1}$ = 0.05, 0.10, 0.15, 0.20, 0.25 and 0.30** (resulting in $\frac{\hat{\beta}_1^2}{p_{exp}} - \frac{\hat{\beta}_2^2}{(1-p_{exp})}$ = 0.015, 0.001, 0.049, 0.166, 0.351 and 0.605).

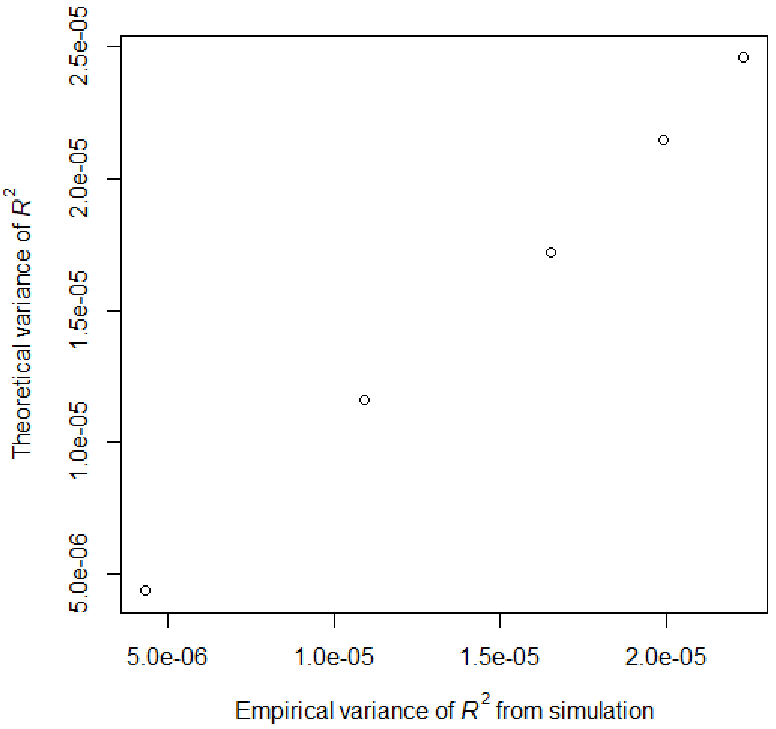


**Supplementary Figure 21: A near-perfect correlation between the theoretical and empirical variances of $R^2$ ($r^2_{y,x_1}$) estimated from 10,000 simulated replicates of ascertained case-control (10000 cases and 10000 controls) assuming 5% disease prevalence and 20000 individuals.** Simulations of $y$, $x_1$ and $x_2$ were based on a correlation structure $\begin{bmatrix} 1 & r_{y,x_1} & r_{y,x_2} \\ r_{y,x_1} & 1 & r_{x_1,x_2} \\ r_{y,x_2} & r_{x_1,x_2} & 1 \end{bmatrix} = \begin{bmatrix} 1 & various & 0.141 \\ various & 1 & 0.800 \\ 0.141 & 0.8 & 1 \end{bmatrix}$ and $R^2$ ($r^2_{y,x_1}$) was obtained from a model $y = x_1 + e$ in each replicate. Following the correlation structure and disease prevalence, we simulated 200,000 dependent and explanatory variables and randomly selected 10000 cases and 10000 controls. The empirical variance of $R^2$ over 10,000 replicates was estimated. The theoretical variance of $R^2$ was obtained from eq. (5). Each data point in the diagonal represents the variance of $R^2$ with $r^2_{y,x_1}$ = 0.02, 0.04, 0.06, 0.08, and 0.1.

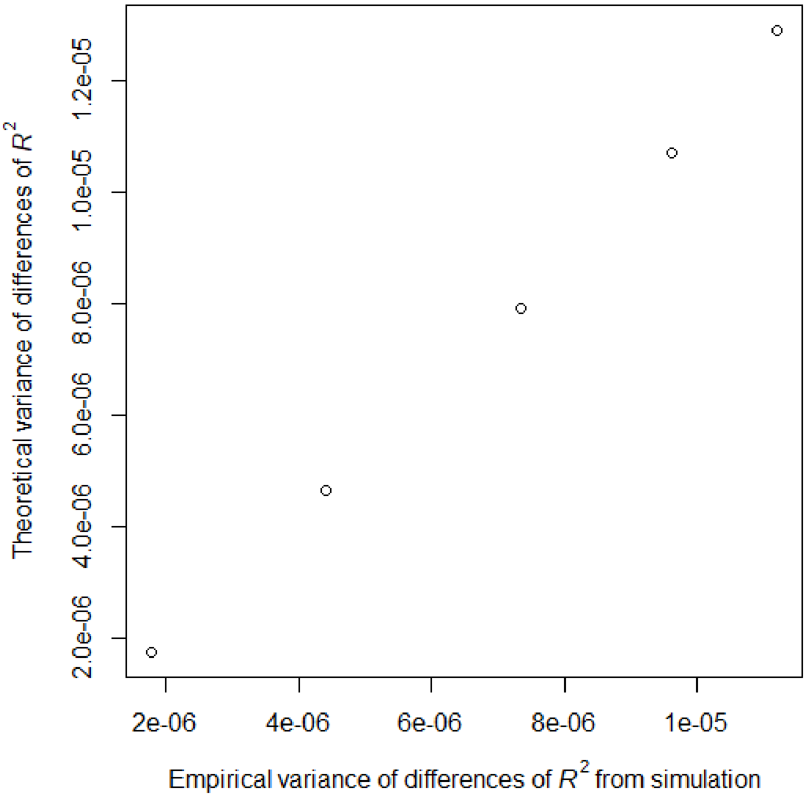


**Supplementary Figure 22: A near-perfect correlation between the theoretical and empirical variances of $R^2$ difference ($r^2_{y,x_1} - r^2_{y,x_2}$) estimated from 10,000 simulated replicates of ascertained case-control (10000 cases and 10000 controls) assuming 5% disease prevalence and 20000 individuals.** Simulations of $y$, $x_1$ and $x_2$ were based on a correlation structure $\begin{bmatrix} 1 & r_{y,x_1} & r_{y,x_2} \\ r_{y,x_1} & 1 & r_{x_1,x_2} \\ r_{y,x_2} & r_{x_1,x_2} & 1 \end{bmatrix} = \begin{bmatrix} 1 & various & 0.141 \\ various & 1 & 0.800 \\ 0.141 & 0.8 & 1 \end{bmatrix}$ and $r^2_{y,x_1}$ and $r^2_{y,x_2}$ were obtained from models $y = x_1 + e$ and $y = x_2 + e$, respectively, to get their difference in each replicate. . Following the correlation structure and disease prevalence, we simulated 200,000 dependent and explanatory variables and randomly selected 10000 cases and 10000 controls. The empirical variance of $R^2$ over 10,000 replicates was estimated. The theoretical variance of $R^2$ was obtained from eq. (5). Each data point in the diagonal represents the variance of $r^2_{y,x_1} - r^2_{y,x_2}$ with $r^2_{y,x_1} - r^2_{y,x_2}$ = 0, 0.02, 0.04, 0.06, and 0.08.

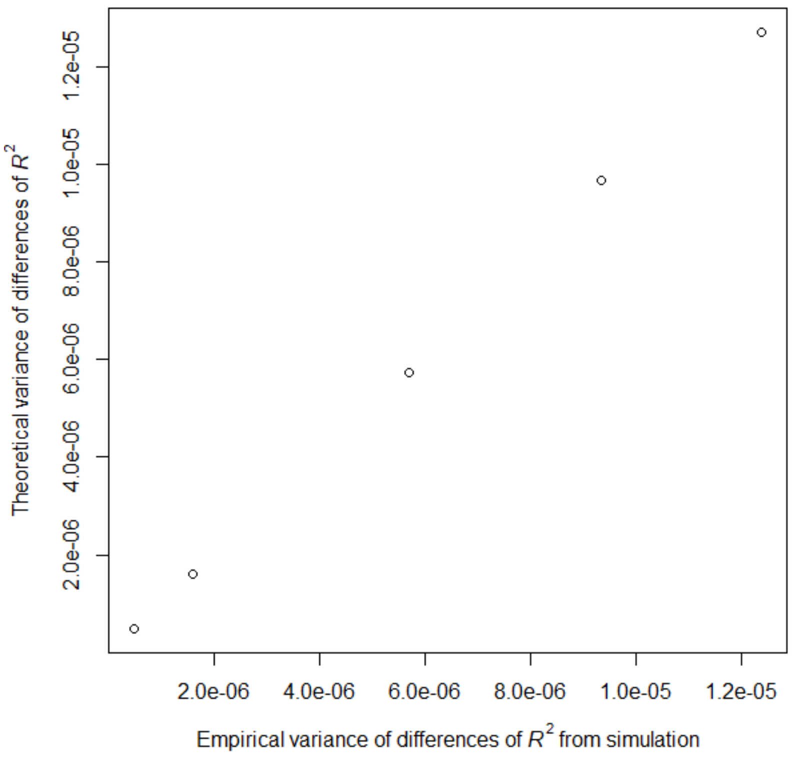


**Supplementary Figure 23: A near-perfect correlation between the theoretical and empirical variances of $R^2$ difference ($r^2_{y,(x_1,x_2)} - r^2_{y,x_1}$) estimated from 10,000 simulated replicates of ascertained case-control (10000 cases and 10000 controls) assuming 5% disease prevalence and 20000 individuals.** Simulations of $y$, $x_1$ and $x_2$ were based on a correlation structure $\begin{bmatrix} 1 & r_{y,x_1} & r_{y,x_2} \\ r_{y,x_1} & 1 & r_{x_1,x_2} \\ r_{y,x_2} & r_{x_1,x_2} & 1 \end{bmatrix} = \begin{bmatrix} 1 & various & 0.141 \\ various & 1 & 0.800 \\ 0.141 & 0.8 & 1 \end{bmatrix}$ and $r^2_{y,(x_1,x_2)}$ and $r^2_{y,x_1}$ were obtained from models $y = x_1 + x_2 + e$ and $y = x_1 + e$, respectively, to get their difference in each replicate. Following the correlation structure and disease prevalence, we simulated 200,000 dependent and explanatory variables and randomly selected 10000 cases and 10000 controls. The empirical variance of $R^2$ over 10,000 replicates was estimated. The theoretical variance of $R^2$ was obtained from eq. (5). Each data point in the diagonal represents the variance of $r^2_{y,x_1} - r^2_{y,x_2}$ with $r^2_{y,x_1} - r^2_{y,x_2}$ = 0, 0.04, 0.08, 0.12, and 0.16.

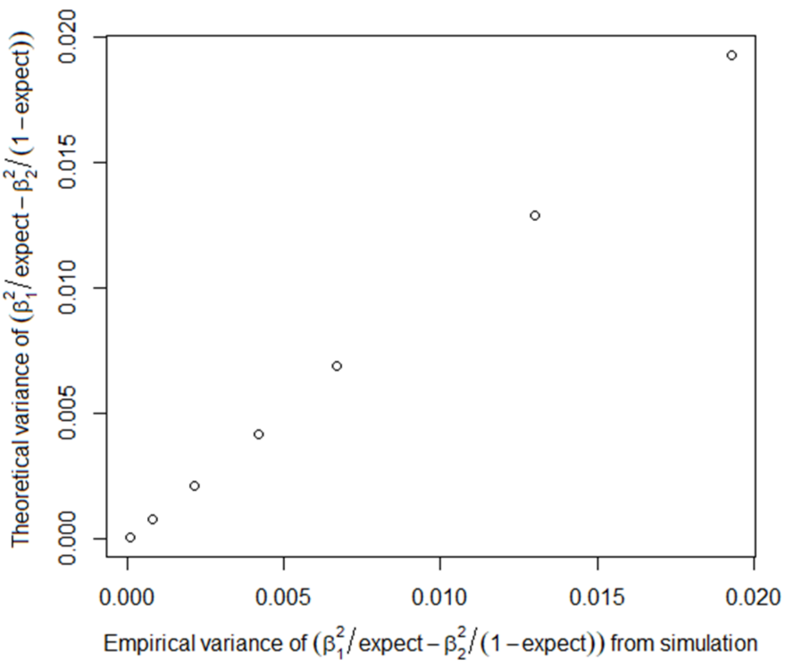


**Supplementary Figure 24: A near-perfect correlation between the theoretical and empirical variances of $\frac{\hat{\beta}_1^2}{p_{exp}} - \frac{\hat{\beta}_2^2}{(1-p_{exp})}$ estimated from 10,000 simulated replicates of ascertained case-control (10000 cases and 10000 controls) assuming 5% disease prevalence and 20000 individuals.** Simulations of $y$, $x_1$ and $x_2$ were based on a correlation structure $\begin{bmatrix} 1 & r_{y,x_1} & r_{y,x_2} \\ r_{y,x_1} & 1 & r_{x_1,x_2} \\ r_{y,x_2} & r_{x_1,x_2} & 1 \end{bmatrix} = \begin{bmatrix} 1 & various & 0.148 \\ various & 1 & 0.610 \\ 0.148 & 0.610 & 1 \end{bmatrix}$, and $\hat{\beta}_1^2$ and $\hat{\beta}_2^2$ were obtained from a multiple regression model $y = x_1 + x_2 + e$ to get the difference of scaled squared regression coefficients in each replicate. It was assumed that the expectation is known ($p_{exp}$ = 0.04 was used). Following the correlation structure and disease prevalence, we simulated 200,000 dependent and explanatory variables and randomly selected 10000 cases and 10000 controls. The empirical variance of $\frac{\hat{\beta}_1^2}{p_{exp}} - \frac{\hat{\beta}_2^2}{(1-p_{exp})}$ over 10,000 replicates was estimated. The theoretical variance of $\frac{\hat{\beta}_1^2}{p_{exp}} - \frac{\hat{\beta}_2^2}{(1-p_{exp})}$ was obtained from eq. (17). Each data point in the diagonal represents the variance of $\frac{\hat{\beta}_1^2}{p_{exp}} - \frac{\hat{\beta}_2^2}{(1-p_{exp})}$ with $r_{y,x_1}$ = 0.05, 0.10, 0.15, 0.20, 0.25 and 0.30 (resulting in $\frac{\hat{\beta}_1^2}{p_{exp}} - \frac{\hat{\beta}_2^2}{(1-p_{exp})}$ = 0.077, -0.015, 0.254, 0.876, 1.819 and 3.090).

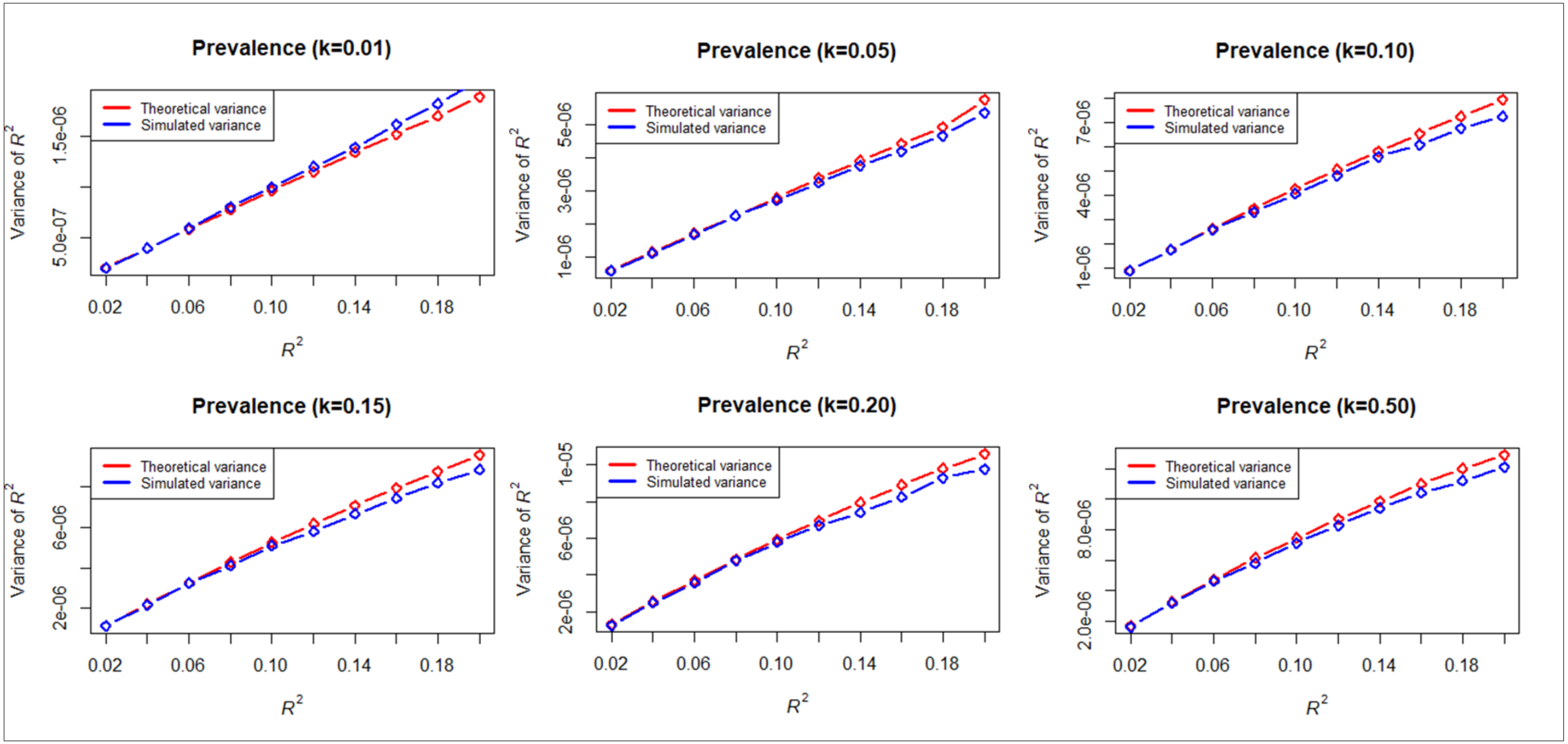


**Supplementary Figure 25: The empirical and theoretical variances become disagreed when $R^2$ values are more than 0.1 for binary responses, noting that $R^2$ > 0.1 is not frequently observed (see Supplementary Table 2).** Simulations of $y$, $x_1$ and $x_2$ were based on a correlation structure $\begin{bmatrix} 1 & r_{y,x_1} & r_{y,x_2} \\ r_{y,x_1} & 1 & r_{x_1,x_2} \\ r_{y,x_2} & r_{x_1,x_2} & 1 \end{bmatrix} = \begin{bmatrix} 1 & various & 0.141 \\ various & 1 & 0.800 \\ 0.141 & 0.8 & 1 \end{bmatrix}$ and $R^2$ ($r^2_{y,x_1}$) was obtained from a model $y = x_1 + e$ in each replicate. Following the correlation structure and disease prevalence, we simulated 30,000 dependent and explanatory variables. The empirical variance of $R^2$ over 10,000 replicates was estimated. The theoretical variance of $R^2$ was obtained from eq. (5). Each data point in the diagonal represents the variance of $R^2$ ranged from 0.02 to 0.2.

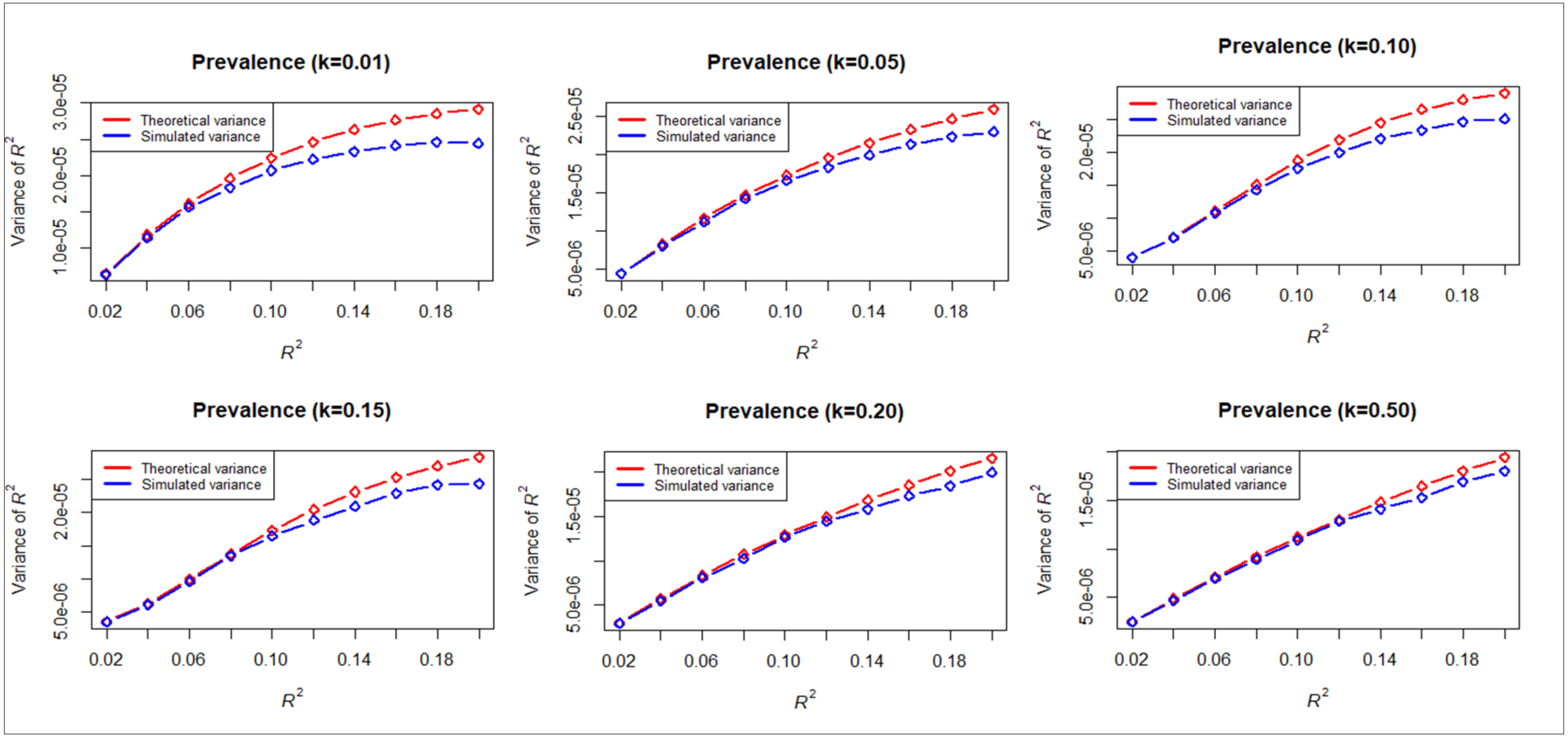


**Supplementary Figure 26: The empirical and theoretical variances become disagreed when $R^2$ values are more than 0.1 for ascertained case-control samples in the reference dataset (10000 cases and 10000 controls), noting that $R^2$ > 0.1 is not frequently observed (see Supplementary Table 2).** Simulations of $y$, $x_1$ and $x_2$ were based on a correlation structure $\begin{bmatrix} 1 & r_{y,x_1} & r_{y,x_2} \\ r_{y,x_1} & 1 & r_{x_1,x_2} \\ r_{y,x_2} & r_{x_1,x_2} & 1 \end{bmatrix} = \begin{bmatrix} 1 & various & 0.141 \\ various & 1 & 0.800 \\ 0.141 & 0.8 & 1 \end{bmatrix}$ and $R^2$ ($r^2_{y,x_1}$) was obtained from a model $y = x_1 + e$ in each replicate. Following the correlation structure and disease prevalence, we simulated 200,000 dependent and explanatory variables and randomly selected 10000 cases and 10000 controls. The empirical variance of $R^2$ over 10,000 replicates was estimated. The theoretical variance of $R^2$ was obtained from eq. (5). Each data point in the diagonal represents the variance of $R^2$ ranged from 0.02 to 0.2.

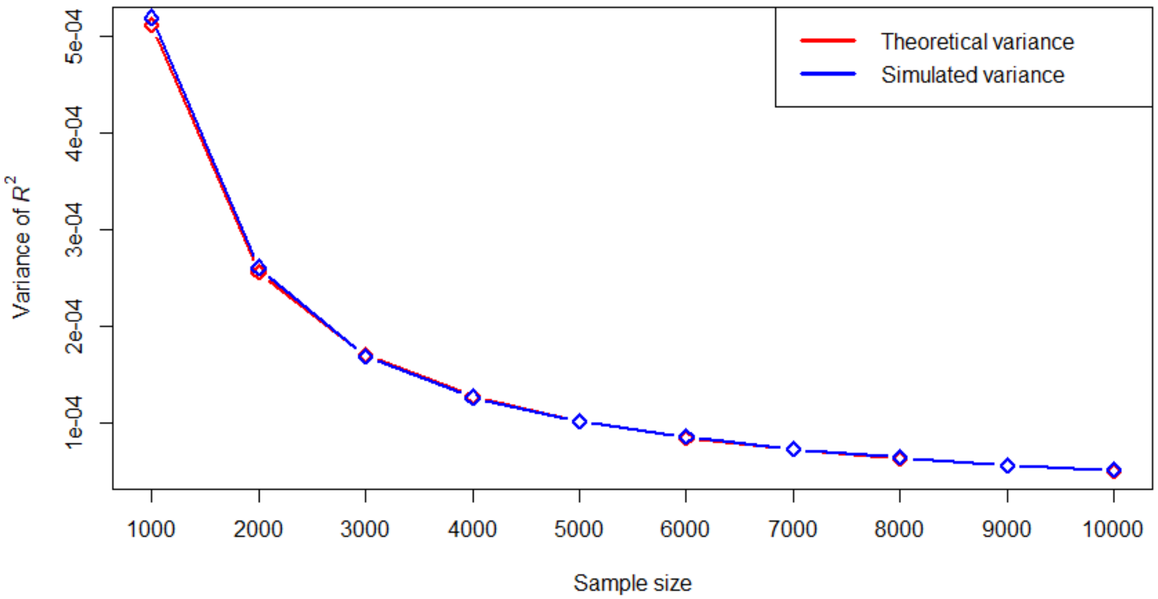


**Supplementary Figure 27: The empirical and theoretical variances is agreed even with sample size 2000 for quantitative phenotypes.** Simulations of $y$, $x_1$ and $x_2$ were based on a correlation structure $\begin{bmatrix} 1 & r_{y,x_1} & r_{y,x_2} \\ r_{y,x_1} & 1 & r_{x_1,x_2} \\ r_{y,x_2} & r_{x_1,x_2} & 1 \end{bmatrix} = \begin{bmatrix} 1 & 0.44 & 0.31 \\ 0.44 & 1 & 0.800 \\ 0.31 & 0.8 & 1 \end{bmatrix}$ and $R^2$ ($r^2_{y,x_1}$) was obtained from a model $y = x_1 + e$ in each replicate. Following the correlation structure and disease prevalence, we simulated dependent and explanatory variables. The empirical variance of $R^2$ over 10,000 replicates was estimated. The theoretical variance of $R^2$ was obtained from eq. (5). Each data point in the diagonal represents the variance of $R^2$ for different sample size.

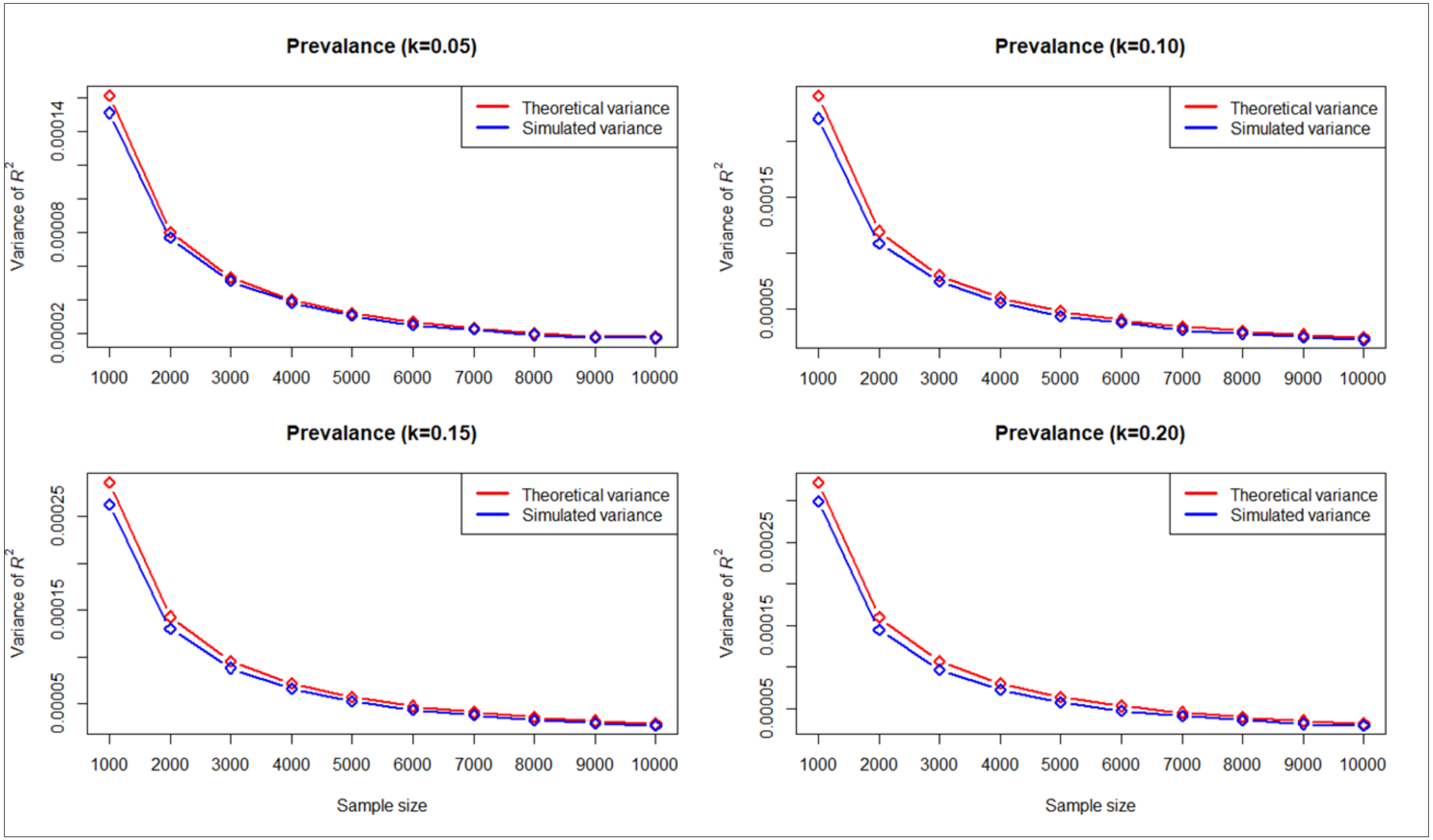


**Supplementary Figure 28: The empirical and theoretical variances become disagreed when sample size is < 5000 for binary responses under scenario of different prevalence rate (k).** Simulations of $y$, $x_1$ and $x_2$ were based on a correlation structure $\begin{bmatrix} 1 & r_{y,x_1} & r_{y,x_2} \\ r_{y,x_1} & 1 & r_{x_1,x_2} \\ r_{y,x_2} & r_{x_1,x_2} & 1 \end{bmatrix} = \begin{bmatrix} 1 & 0.44 & 0.31 \\ 0.44 & 1 & 0.800 \\ 0.31 & 0.8 & 1 \end{bmatrix}$ and $R^2$ ($r^2_{y,x_1}$) was obtained from a model $y = x_1 + e$ in each replicate. Following the correlation structure and disease prevalence, we simulated dependent and explanatory variables. The empirical variance of $R^2$ over 10,000 replicates was estimated. The theoretical variance of $R^2$ was obtained from eq. (5). Each data point in the diagonal represents the variance of $R^2$ for different sample size.

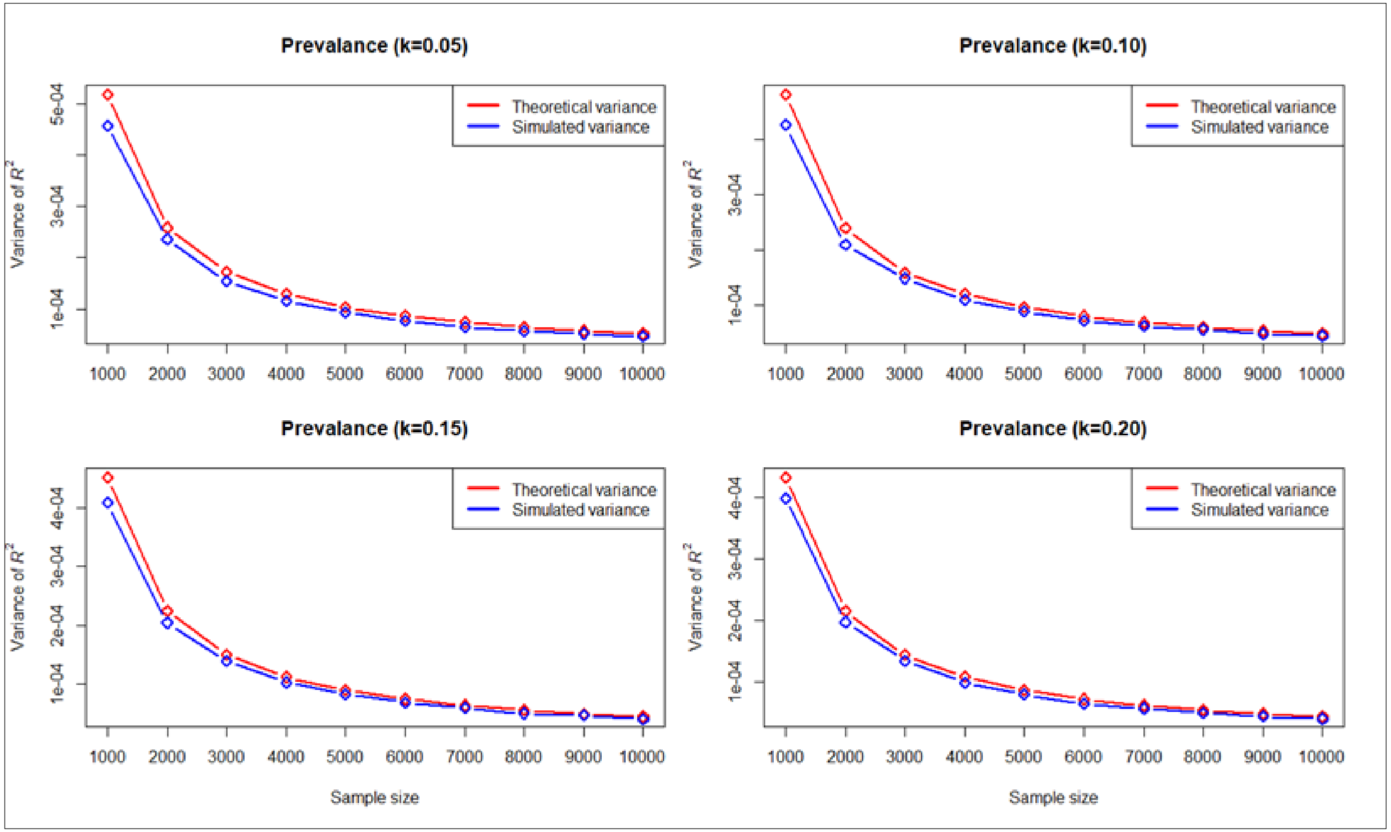


**Supplementary Figure 29: The empirical and theoretical variances become disagreed when sample size is < 5000 for ascertained case-control samples in the reference dataset (50% cases and 50% controls) under scenario of different prevalence rate (k).** Simulations of $y$, $x_1$ and $x_2$ were based on a correlation structure $\begin{bmatrix} 1 & r_{y,x_1} & r_{y,x_2} \\ r_{y,x_1} & 1 & r_{x_1,x_2} \\ r_{y,x_2} & r_{x_1,x_2} & 1 \end{bmatrix} = \begin{bmatrix} 1 & 0.44 & 0.31 \\ 0.44 & 1 & 0.800 \\ 0.31 & 0.8 & 1 \end{bmatrix}$ and $R^2$ ($r^2_{y,x_1}$) was obtained from a model $y = x_1 + e$ in each replicate. Following the correlation structure and disease prevalence, we simulated 100,000 dependent and explanatory variables and randomly selected cases and controls. The empirical variance of $R^2$ over 10,000 replicates was estimated. The theoretical variance of $R^2$ was obtained from eq. (5). Each data point in the diagonal represents the variance of $R^2$ for different sample size.

**Supplementary Table 1:** Number of SNPs across different *P*-value thresholds for BMI and cholesterol for UKBB and BBJ

| ***P*-value Threshold** | **BMI** | | **Cholesterol** | |
|---|---|---|---|---|
| | **No of SNPs (UKBB)** | **No of SNPs (BBJ)** | **No of SNPs (UKBB)** | **No of SNPs (BBJ)** |
| 1 | 4113630 | 4113630 | 4113630 | 4113630 |
| 0.5 | 2539432 | 2365077 | 2254467 | 2143406 |
| 0.4 | 2199702 | 1996741 | 1864917 | 1746560 |
| 0.3 | 1841948 | 1610857 | 1468402 | 1346257 |
| 0.2 | 1442727 | 1201525 | 1059630 | 936383 |
| 0.1 | 976865 | 746948 | 618466 | 508651 |
| 5e-02 | 675502 | 475337 | 376346 | 280526 |
| 1e-02 | 318902 | 128704 | 140757 | 76284 |
| 1e-03 | 134943 | 57272 | 54829 | 19216 |
| 1e-04 | 67528 | 24320 | 30741 | 8636 |

**Supplementary Table 2:** The AUC values (reported in Khera et al.[1] ) and $R^2$ values converted from the AUC given the sample size, prevalence in discovery and testing datasets.

| **Disease** | **Prevalence in discovery GWAS (*n*)** | **Prevalence in validation dataset** | **AUC (95% CI) in validation dataset** | **Predictive ability ($R^2$)** |
|---|---|---|---|---|
| CAD | 60,801 cases and 123,504 controls (32.9%)[2] | 3,963 cases and 116,317 controls (3.4%) | 0.81 (0.80–0.81) | 0.040 |
| Atrial fibrillation | 17,931 cases and 115,142 controls (13.4%)[3] | 2,024 cases and 118,256 controls (1.7%) | 0.77 (0.76–0.78) | 0.016 |
| Type 2 diabetes | 6,676 cases and 132,532 controls (16.7%)[4] | 2,785 cases and 117,495 controls (2.4%) | 0.72 (0.72–0.73) | 0.012 |
| Inflammatory bowel disease | 2,882 cases and 21,770 controls (37.2)[5] | 1,360 cases and 118,920 controls (1.1%) | 0.63 (0.62–0.65) | 0.003 |
| Breast cancer | 122,977 cases and 105,974 controls (53.7)[6] | 2,576 cases and 60,771 controls (4.1%) | 0.68 (0.67–0.69) | 0.017 |

These results were quoted from Khera et al.[1] $R^2$ values were converted from the AUC using the well-established theory[7; 8].
